# Interspecies Differential Gene Expression Analysis with Regularized Phylogenetic Linear Models

**DOI:** 10.64898/2026.06.30.734542

**Authors:** Mélina Gallopin, Maëlle Daunesse, Olivier Lespinet, Arnaud Liehrmann, Paul Bastide

## Abstract

Comparative transcriptomic datasets are increasingly used to investigate the molecular basis of phenotypic diversification across species. However, finding genes that are differentially expressed (DE) between lineages remains challenging, for two main reasons. First, the random evolutionary drift can blur the signal left by lineage-specific shifts in mean expression, and induces phylogenetic correlations that, if ignored, can widely inflate the False Discovery Rate (FDR), i.e., the amount of spuriously detected genes. Second, DE analysis from RNA-Seq data involves multiple testing on many genes for a small number of individual measurements with high noise, and requires dedicated statistical tools. Traditional DE tools, such as limma, and classical Phylogenetic Comparative Methods (PCMs), such as the Expression Variance and Evolution (EVE) model, are both designed to tackle one of these two challenges alone, but both fail in the context of inter-species RNA-Seq data. In this work, we present phyloDE, a new tool for inter-species DE, that aims at taking the best from both approaches. On simulations based on a recently published four-species rodent dataset, we show that, contrary to other methods, phyloDE correctly controls the FDR in all settings, while keeping a reasonable power. When reanalyzing the empirical dataset, phyloDE discovers more DE genes that exhibit consistent changes in their cis-regulatory landscape compared to EVE in all the experimental settings. The method is implemented in R, with an interface inheriting from limma.

## Introduction

Shifts in gene expression across species can be the signature of important evolutionary mechanisms, that may drive the observed phenotypic divergence in many groups of organisms (Romero *et al*., 2012; Price *et al*., 2022). In recent years, interspecies transcriptomic datasets have become more common, and have been used to detect differentially expressed (DE) genes on nested lineages in numerous families of species, including amniotes (Brawand *et al*., 2011), primates (Perry *et al*., 2012), rodents (Chalopin *et al*., 2026), crayfishes (Stern and Crandall, 2018) and catfishes (Phelps *et al*., 2026), among others (Torres-Oliva *et al*., 2016; Berthelot *et al*., 2018; Catalán *et al*., 2019; El Taher *et al*., 2021).

One of the main challenges when studying such interspecies datasets is to distinguish meaningful expression shifts from both technical and phylogenetic noise (Bastide *et al*., 2023). RNA-Seq data usually involves a small number of noisy individual measurements for a large number of genes. Differential expression analysis in this context requires dedicated statistical methods, that are designed to exploit information from the entire dataset to retain some statistical power while controlling for the false discovery rate (FDR), i.e., the amount of spuriously detected neutral genes. These methods, implemented, e.g., in the R packages limma, DESeq2 or edgeR (Anders and Huber, 2010; Robinson and Oshlack, 2010; Love *et al*., 2014; Ritchie *et al*., 2015), are widely used in the classical, single species design. They rely on sharing information between genes, instead of considering each of them independently. In particular, limma uses an empirical Bayes procedure, that assumes a common prior for all the variance parameters for all the gene expressions. This prior on the variance can be made a fonction of the mean gene expression, in order to account for the mean-variance relationship that is often exhibited by RNA-Seq datasets (limma-trend, Law *et al*., 2014). It then leverages this hierarchical structure in a three steps procedure, first estimating all the variance parameters independently; then estimating the prior from these raw estimates using a robust fitting method (Phipson *et al*., 2016); and finally *regularizing* the variance estimates by replacing them with their empirical posterior estimator, that integrates information from all the genes on each individual expression. This produces *moderated t*-tests, that are shown to be more powerful than simple *t*-tests (Smyth, 2004).

However, phylogenetic inertia, induced by neutral evolution or random drift, can produce a signal in the data that, if ignored, may be wrongly interpreted as differential expression. This problem can be mitigated using well-though data-collection designs. For instance, Parker *et al*. (2019) study gene expression shift in sexual versus asexual stick insects with edgeR, using a paired design that associate each sexual species with a closely related asexual one. However, most datasets present an unbalanced design, comparing one monophyletic group of species to others, as, e.g., primates versus other mammals (Chen *et al*., 2019). Traditional methods, that assume that all the measurements are independent, are no longer suitable in this context. In the context of within array replicate spots in microarray experiments, Smyth *et al*. (2005) extended the limma frameworks to measurements that are *block-structured*, implemented in the R function duplicatedCorrelation. It assumes that observations within a given block are correlated with a single correlation parameter, while observations between blocks are independent. It then strongly regularizes all the correlation parameters across genes, by assuming that all genes share the same correlation parameter. In the context of inter-species measurements, this would be equivalent to assuming that replicates within each species are correlated, but that species themselves are independent (Bastide *et al*., 2023), that is, replacing the true phylogenetic tree by a star tree. This block correlation structure associated with star trees is illustrated on the bottom row of Figure 3. Some extensions of these methods, based on a linear mixed model (Sun *et al*., 2017; Yu *et al*., 2019; Hoffman and Roussos, 2020) aim at taking into account non-independent measurements, including complex structures such as kinship matrices (Dill-McFarland *et al*., 2023), that can be linked with phylogenetic variance-covariance matrices (Bastide *et al*., 2018; Schraiber *et al*., 2024). However, these methods are not designed to model gene expression along a phylogenetic tree, and they cannot fit models of adaptive or stabilizing selection such as the Ornstein-Uhlenbeck (OU) process (Hansen, 1997), that is widely used in phylogenetic comparative methods (Cooper *et al*.,2016), and that has been shown to be well suited to model gene expression evolution on a wide range of datasets (Dimayacyac *et al*., 2023).

Phylogenetic comparative methods (PCMs) precisely aim at taking into account the shared phylogenetic histories of individuals when analyzing continuous or discrete measurements on present-day species (Felsenstein, 2004; Harmon *et al*., 2010; Revell and Harmon, 2022). Popular all-purpose R implementations include, among others, geiger (Pennell *et al*., 2014), phylolm Ho and Ané (2014a) and mvMORPH Clavel *et al*. (2015). In the context of gene expression analysis, Rohlfs *et al*. (2014) proposed the Expression Variance and Evolution method (EVE), that relies on a OU model for trait evolution, with additional within-species variance. In particular, the twoThetaTest method implemented in the R package evemodel (Gillard *et al*., 2021, referred to as the eve-ttt method for short) uses a phylogenetic ANOVA (Garland *et al*., 1993; Rohlfs and Nielsen, 2015) to detect DE genes across specified lineages. The EVE method pioneered the use of PCMs in interspecies differential gene expression analysis, and has been widely adopted in the community (Chen *et al*., 2019; Gillard *et al*., 2021; Daunesse *et al*., 2026; Chalopin *et al*., 2026; Phelps *et al*., 2026). However, just like the classical ANOVA was shown to be ill-suited for classical RNA-Seq analysis (Jeanmougin *et al*., 2010), the standard phylogenetic ANOVA of the EVE method has some limitations. Importantly, it performs gene-by-gene tests independently, and does not exploit any mechanism to share information between genes. As recalled above, empirical Bayes approaches are an example of such a mechanism, that have been highly successful in classical RNA-Seq methods (Smyth, 2004), yielding increased statistical power, especially for datasets with a small number of individual measurements.

Regularization, including empirical Bayes, has been applied in a phylogenetic context to estimate the evolutionary rate (Bertram *et al*., 2022) or the selection strength (Gu *et al*., 2019; Yang *et al*., 2019). In this work, the precise estimation of the variance parameters of the model is not our main concern, as our primary objective is differential expression analysis, i.e., the detection of genes that exhibit a difference in mean expression between identified groups of species. As in Rohlfs and Nielsen (2015), such differences can be interpreted as representing shifts in the optimal expression value between groups of species, and hence be a marker of selection. We introduce phyloDE, a method specifically designed for interspecies differential expression analysis. It relies on both PCMs and empirical Bayes regularization to correctly account for the evolutionary structure and the technical noise of RNA-Seq data collected on several species. Just as limma and evemodel, our method takes as input a table quantifying the expression levels of one-to-one orthologous genes, and assumes that the data has been properly normalized before-hand, using a metric that takes the gene length into account (see Bastide *et al*., 2023 for a discussion of the choice of normalization factors in this context). Using the statistical framework from Bastide *et al*. (2023) and implemented in compcodeR (Soneson, 2014), we simulated synthetic data based on a recent RNA-seq dataset (Daunesse *et al*., 2026), a dataset originally generated to study the coordinated changes in gene expression and regulatory activity in heart and liver tissues in African mole-rats. Among all the datasets previously analyzed with evemodel, it has one of the least favorable design, with only 4 rodent species included in the analysis. Such a small dataset proved challenging to existing methods, that had very little power (eve-ttt) or high rates of false discoveries (limma) on simulated data with known ground truth. By contrast, we show that the phyloDE method increased the power of detection of DE genes between species, while controlling the FDR. When applied to the original empirical dataset of Daunesse *et al*. (2026), phyloDE increased the number of DE genes compared to eve-ttt, while finding genes that exhibited consistent changes in their cis-regulatory landscape in all the experimental settings. The phyloDE method is implemented in an R package freely available on GitHub, and has been designed to have a syntax similar to the familiar software limma, while relying on the fast PCM algorithm of phylolm.

## Results

### The phyloDE method

Any inter-species differential gene expression analysis from RNA-Seq data must face four main challenges: (i) inter-species measurements are not independent, and the method must account for the evolutionary processes that shaped the distribution of the data; (ii) some of the parameters of the evolutionary model are difficult to estimate from present-day data only, and a careful parametrization and regularization must be used; (iii) differential expression analysis usually considers several thousands of genes for a limited number of individuals, and moderation techniques that leverage the entire dataset when testing each individual genes can improve the statistical power substantially; and (iv) RNA-Seq count data can have specific noise structures including, e.g., a possible mean-variance relationship, that need to be taken into account.

At its core, the phyloDE method combines the strengths of phylogenetic linear regression and empirical Bayes regularization to address those four challenges. phyloDE relies on the standard PCM tool phylolm (Ho and Ané, 2014a) to perform phylogenetic regression (challenge (i)). phyloDE uses a careful parametrization of the Ornstein-Uhlenbeck process through biologically meaningful quantities (Cornuault, 2022), that can be regularized accros genes using an *ad-hoc* extension of the duplicatedCorrelation method of Smyth *et al*. (2005) from a block-structure (star tree, see Introduction) to a phylogenetic structure (challenge (ii)). These regularized parameters then unlock the use of the limma framework (Ritchie *et al*., 2015) on the de-correlated residuals, that includes robust empirical Bayes moderated *t*-tests (Smyth, 2004; Phipson *et al*., 2016) (challenge (iii)) and a mean-variance relationship correction using the limma-trend approach (Law *et al*., 2014), which is known to be particularly important for RNA-Seq data (challenge (iv)). Compared to state-of-the-art methods such as the twoThetaTest method from EVE (Rohlfs *et al*., 2014, referred to as the eve-ttt method for short) and limma with block structured correlation (Smyth *et al*., 2005; Ritchie *et al*., 2015, referred to as limma-cor), phyloDE is the only method to our knowledge that addresses all of the four challenges described above (see Table 1 for a synthetic comparison). The details of our approach and its similarities and differences with previous methods are described in the Material and Method Section.

**Table 1.** Synthetic comparison of phyloDE with the twoThetaTest method from EVE (eve-ttt, Rohlfs *et al*., 2014) and phylolm (Ho and Ané, 2014a), that takes the phylogeny into account but treat each of the genes independently; and limma-cor (Smyth *et al*., 2005), that exploits the regularization accros genes to gain statistical power and deal with RNA-Seq specific technical features but ignores the evolutionary process.

|  | eve-ttt | phylolm | limma-cor | phyloDE |
|---|---|---|---|---|
| (i) Accounts for phylogenetic tree | **yes** | **yes** | no | **yes** |
| (ii) Regularized correlation parameters across genes | no | no | **yes** | **yes** |
| (iii) Empirical Bayes moderated tests | no | no | **yes** | **yes** |
| (iv) Accounts for mean-variance relationships | no | no | **yes** | **yes** |

### Simulation Study

Our main simulation study relies on an empirical dataset extracted from Daunesse *et al*. (2026), who collected RNA-Seq data concerning gene expression of four species: the naked mole-rat (*Heterocephalus glaber*, Hgla), the Damaraland mole-rat (*Fukomys damarensis*, Fdam), the guinea pig (*Cavia porcellus*, Cpor), and the mouse (*Mus musculus*, Mmus). In all simulations, we used a dated phylogenetic tree extracted from the larger mammals tree (Hassler *et al*., 2020), and 3 replicates per species (the minimum number of biological replicates per species that could realistically be sequenced in an experimental setting). As in Daunesse *et al*. (2026), we then considered two different designs for differential expression analysis. The “Hgla” design isolates the naked mole rat from the three other species, while the “MR” design groups the two mole rat species in a single group (see, e.g., Fig. 2, top-row for a schematic view of the tree and designs). Note that both designs are structured in “blocks”, i.e., are consistent with clades on the tree, so that we expect that the phylogenetic signal had a strong impact on differential expression analysis (Bastide *et al*., 2023).

#### Variance parameter estimation is highly variable across genes

To assess the amount of signal available in a typical dataset to estimate the variance parameters, we first simulated a single dataset with 10 000 genes according to a OU process with fixed parameters on the mole rat tree, with no group effect, so that all the species had the same expected trait value, and all the genes had the same variance structure. We then applied evemodel, phylolm and phyloDE on this dataset, and compared the estimated parameters with the true values used for simulation. As the simulated model is the same as the fitting model for all methods, we expected the parameters to be correctly inferred.

Fig. 1 shows the values estimated by evemodel, using the OU parametrization from Rohlfs and Nielsen (2015) (see Material and Methods). The expectation parameter *θ* was correctly estimated, with a mean value across genes close to the true value of 1, and a uni-modal distribution with standard deviation around 0.5. However, the two parameters that control the variance structure across samples, namely, *β* the ratio between the evolutionary variance and the within-population variance and *α* the selection strength, were poorly estimated and highly variable across genes. The distribution of estimated *β* values is bimodal, and ranges between 1.54 *·* 10^*−*2^ and 5.36 *·* 10^9^ (true value is 0.31), while the distribution of estimated *α* values is erratic, ranging between 1.08 and 6.14*·*10^8^ (true value is 1.39). Similar results were obtained on the same example with phylolm and phyloDE (before regularization, see Supplementary Material Section S3). These methods use the same model as EVE, so that the optimized likelihood of the fitted model on each gene was very similar between methods. However, estimations of the parameters were highly variable between methods (see Supplementary Figure S8).

**Figure 1.**
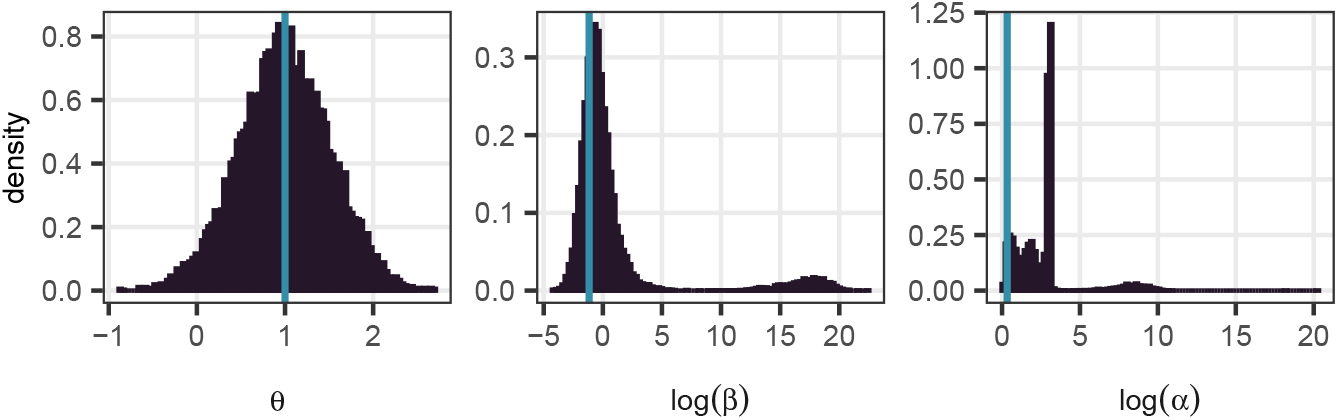
Histogram of estimated parameters using EVE (fonction fitOneTheta from package evemodel, Gillard *et al*., 2021), for 10 000 genes, all simulated with a OU process with fixed parameter values (light blue lines) on the mole rats tree with the three replicates per species. The OU is parametrized as in Rohlfs and Nielsen (2015), with *θ* the trait expected value (true value of 1), *α* the selection strength (true value of 1.39); and *β* the ratio between the evolutionary variance and the within-population variance (true value of 0.31). The variance parameters *α* and *β* are poorly estimated and highly variable across genes (log-scale used in the histograms).

This illustrates the fact that variance parameters such as the selection strength of the OU or the within-species variance are difficult to estimate on small trees. When our goal is differential expression analysis (that is, finding shifts in the mean), we show below that these parameters can be seen as “nuisance parameters” and regularized out for improved DE detection. In this setting, the regularized estimates of the parameters found by phyloDE were close to the true values, so that the model approximately captured the correct correlation structures (see Supplementary Figure S6).

#### phyloDE controls the FDR for realistic count data

Following the procedure of Bastide *et al*. (2023), we used the R package compcodeR (Soneson, 2014) with parameters inferred from the liver raw RNA-Seq data of Daunesse *et al*. (2026) to generate realistic count data, using either the Brownian Motion (BM; Felsenstein, 1985) or the Ornstein-Uhlenbeck (OU; Hansen and Martins, 1996) as the underlying generating processes. We then analyzed the simulated datasets using limma-cor, phylolm, eve-ttt and phyloDE (see Material and Methods for a complete description of the setting). Fig. 2 shows that phyloDE controlled the FDR for every simulation process (BM or OU) and every design, while keeping a reasonable True Positive Rate (TPR). Method limma-cor, that ignores the phylogenetic structure, failed to control the FDR, particularly in the MR design case. eve-ttt and phylolm methods, that both take the phylogenetic structure into account but do not use any regularization failed in two different ways. While phylolm had a very high FDR (more than 50% in all simulation designs), eve-ttt had a very low TPR, particularly in the Hgla design.

**Figure 2.**
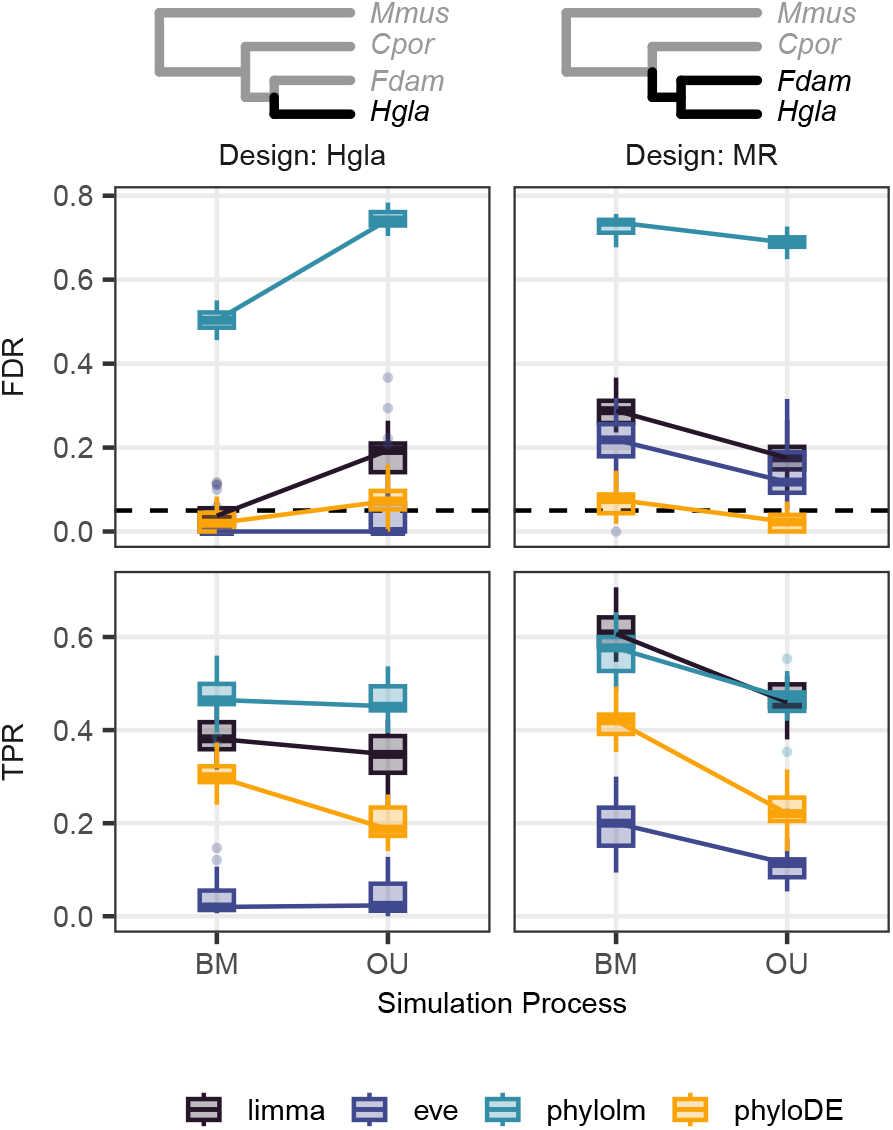
False Discovery Rate (FDR, top row) and True Positive Rate (TPR, bottom row) using Benjamini-Hochberg corrected p-values with a threshold of 5% (dashed horizontal line) for differential expression analysis between species *Hgla* versus the rest of the tree (Design: Hgla, left column), or both mole rat species versus the rest of the tree (Design: MR, right column), for 20 datasets simulated with either the BM or the OU process using compcodeR on the empirical tree with 3 replicates per species, fixed empirical parameters, and a base effect size of 4. Methods are limma-cor (with block correlation and empirical Bayes correction of fitted variances with trend), eve-ttt (with empirical p-values computed using 20 000 simulations under the null), phylolm (assuming a OU process with a fixed root), and phyloDE (assuming a OU process with a fixed root and empirical Bayes correction of fitted variances with trend). Our method phyloDE controlled the FDR, while finding more than 20% of the truly differentially expressed genes.

#### phyloDE leverages the mean-variance relationship for statistical power

The mean-variance relationship is an important feature of RNA-Seq data (challenge (iv) above). phyloDE benefits from the limma implementation of the robust empirical Bayes correction (Phipson *et al*., 2016), which can include a trend in the variance prior. This trend is an alternative method to voom to take the mean-variance relationship into account (Law *et al*., 2014). In our setting, adding a trend significantly increased the statistical power compared to the empirical Bayes correction without trend, with better TPR for a similar FDR control (see Supplementary Figure S4).

#### phyloDE parameters are interpretable

Our method uses a parametrization that is different from that of Rohlfs and Nielsen (2015) and that clearly separates the total variance parameter *σ*^2^ from the correlation structure parameters *λ* and *ρ* (see Equations (4-6) from the Material and Methods). The interpretation of these parameters relies on an equivalence with a Brownian Motion on a transformed tree. A simple BM induces correlations between samples that are directly encoded in the tree shape: the correlation between two samples is equal to their time of shared evolution, i.e., the distance between the root and their most recent common ancestor (Felsenstein, 1985, see Fig. 3 and Eq. (S4) in the Supplementary Material). For instance, according to this BM model, two samples from the same species are weakly correlated for the top left tree of Fig. 3, but highly correlated for the top right tree. A OU process on a tree induces a different correlation structure, that depends on the parameters, and that is not straightforward to directly read from the tree. However, given some parameter values, it is possible to find a transformed tree such that the OU correlation structure from the original tree is equal to the BM correlation on the transformed tree (Blomberg *et al*., 2003; Ho and Ané, 2014a). In our OU parametrisation, *λ* and *ρ* both vary between 0 and 1, and capture different aspects of the correlation structure between samples and between species, that can be graphically understood from the transformed tree. Parameter *λ* is similar to Pagel’s lambda (Pagel, 1999) and controls correlations at the *sample* level: small values mean that the samples within each species are only weakly correlated, while large values mean that all the samples from a same species are highly correlated. Parameter *ρ* was introduced by Cornuault (2022) and controls the correlations at the *species* level: small values mean that the species are structured by the underlying phylogenetic tree, while large values that the species are almost independent from one another. As illustrated in Fig. 3, these two levels can act independently: a trait can be inferred with a small *λ* and a small *ρ*, in which cases species are highly structured but samples within species are almost independent (top left tree), or with a small *λ* and a large *ρ*, so that both species and samples are weakly correlated (bottom left tree). Note that, contrarily to the usual transformation of Ho and Ané (2014a), all the transformed trees have a constant unit height, so that *λ* and *ρ* actually control the correlation structure, and have no impact on the total variance, that is captured by parameter *σ*^2^. We argue that these three parameters *σ*^2^, *λ* and *ρ* capture three separate aspects of the variance and the correlations, which intuitively explains why they are well adapted to the regularization procedures proposed in phyloDE.

**Figure 3.**
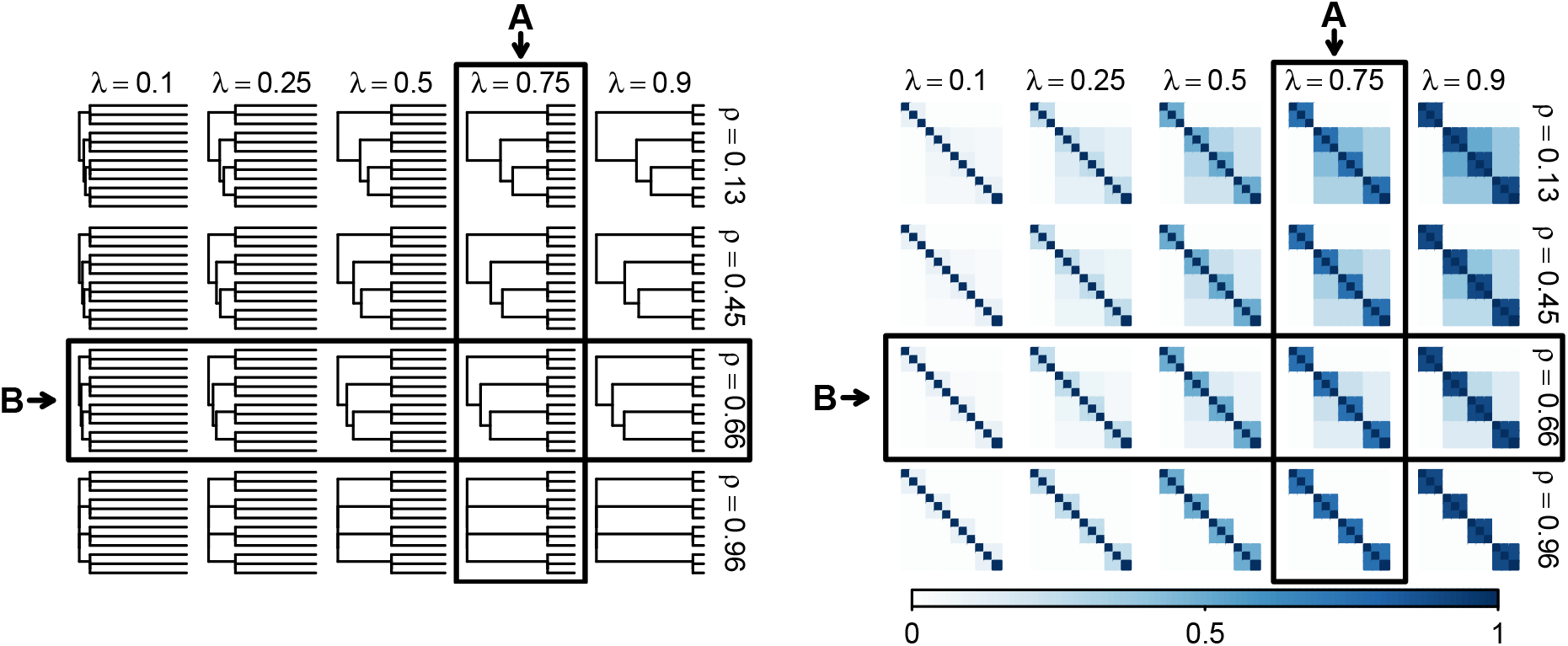
Effects of the parameters on the correlation structure, for the mole rat tree with three replicates per species. Left panel: transformed trees. Right panel: associated correlation matrices. A BM on each of the transformed trees (left panel) has the same correlation matrix (right panel) than a OU with parameters *λ* and *ρ* (see Equations (4-6) from the Material and Methods). Increasing *λ* (from left to right) implies that the samples within each species are more correlated. Increasing *ρ* (from top to bottom) implies that the species are more independent. When *ρ* is close to 1 (bottom row), the tree converges to a star tree, and the correlation matrix is block-structured, similar to the structured assumed by limma-cor. Both transformations in *λ* and *ρ* have distinct effects on the correlation structure, and can be varied independently. Squared column **A** and row **B** correspond to parameters varying in matching panels of Fig. 4.

#### phyloDE controls the FDR for a wide range of parameters

To explore the behavior of the methods for a larger range of parameters, we reproduced the simulations above using direct Gaussian simulations from the OU. As they do not include a count data generation layer, these simulations use the same model for simulation and inference for the three methods based on phylogenetic regression (phyloDE, phylolm and eve-ttt). Again, we see on Figure 4 that phyloDE controlled the FDR for a wide range of parameters, ranging from tree-structured (small *ρ* values) to independent species (large *ρ* values, left panel), and from independent replicates (small *λ* values) to species-structured (large *λ* values, right panel). The MR design led to particularly inflated FDR for other methods. Strong phylogenetic signal (large values of *λ*) caused FDR found by phylolm to sharply increase and TPR found by eve-ttt to drop.

**Figure 4.**
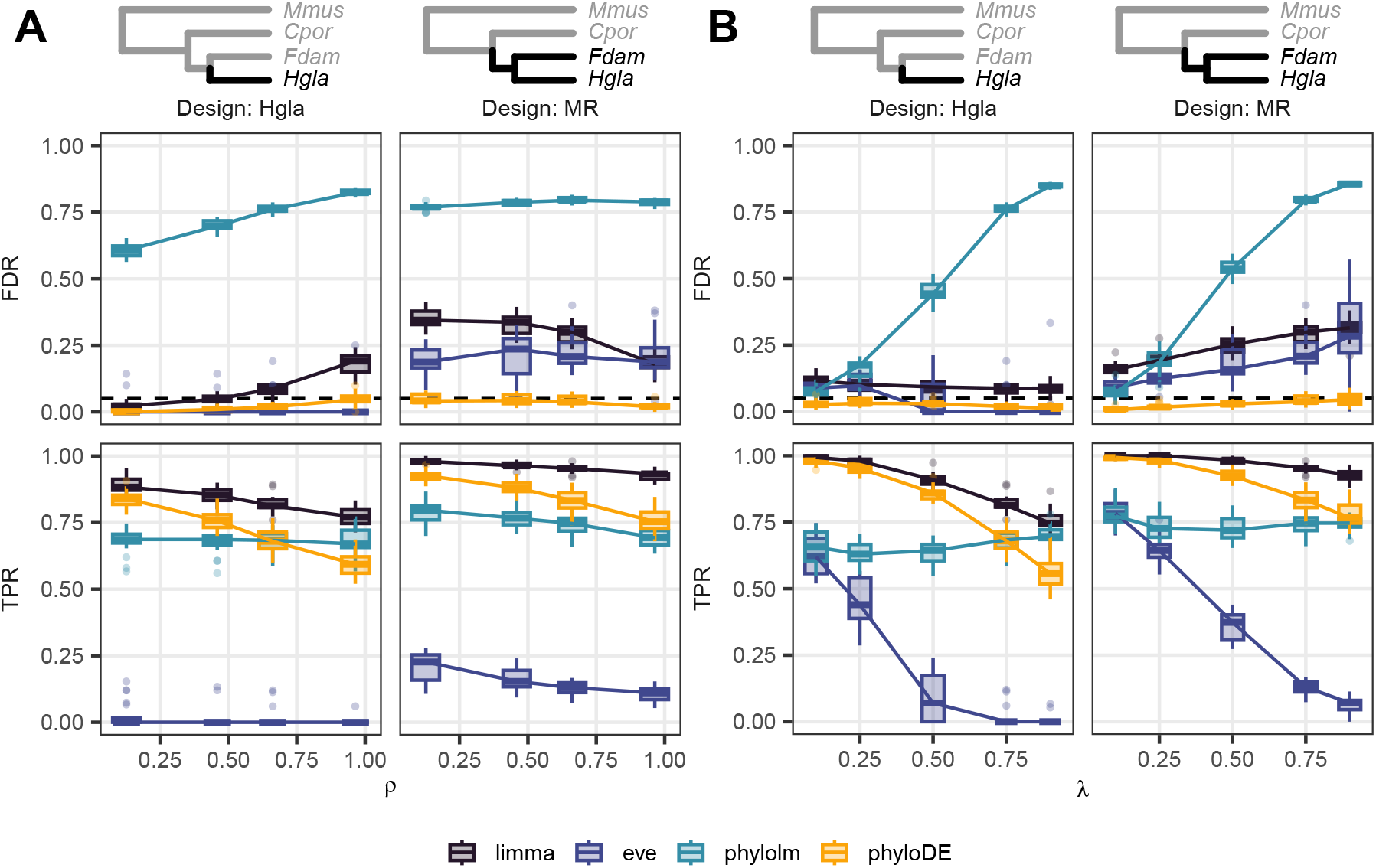
False Discovery Rate (FDR, top row) and True Positive Rate (TPR, bottom row) using Benjamini-Hochberg corrected p-values with a threshold of 5% (dashed horizontal line) for differential expression analysis between species *Hgla* versus the rest of the tree (Design: Hgla), or both mole rat species versus the rest of the tree (Design: MR), for 20 datasets simulated the OU process on the empirical tree with 3 replicates per species, with unit variance and a base effect size of 4. **A**: *λ* is fixed to 0.75, while *ρ* varies (see squared column in Fig. 3). **B**: *ρ* is fixed to 0.66, while *λ* varies (see squared row in Fig. 3). Methods are limma (with block correlation and empirical Bayes correction of fitted variances with trend), eve-ttt (with empirical p-values computed using 20 000 simulations under the null), phylolm (assuming a OU process with a fixed root), and phyloDE (assuming a OU process with a fixed root and empirical Bayes correction of fitted variances with trend). Our method phyloDE controlled the FDR, while finding more than 50% of the truly differentially expressed genes.

#### phyloDE is robust to simulation parameters varying across genes

To regularize the parameters across genes (challenge (ii) above), phyloDE makes the strong assumption that all the genes share the same correlation structure. To challenge this hypothesis, we generated count datasets in the compcodeR framework with phylogenetic parameters varying across all the genes, either with a low or high variance (see Supplementary Figures S2 and S3). This experiment showed that phyloDE was robust to a violation of the common structure assumption, as it controlled the FDR even in the high variance setting, and had a consistent TPR. The other phylogenetic methods phylolm and eve-ttt, that do not rely on this assumption, did not improve when the assumption was relaxed and still exhibited, respectively, a very high FDR or a very low TPR.

#### phyloDE results scale to larger phylogenetic trees

We simulated RNA-Seq count data on trees with 4 to 17 species, extracted from the Crayfish (Stern and Crandall, 2018) and Mammal (Chen *et al*., 2019) datasets, and tested the ability of phyloDE to detect lineage specific mean expression shifts. While the results of all methods improved with the dataset size, phyloDE remained the method with the lowest FDR across all designs (Figures S10 and S11). Both limma-cor and phylolm still exhibited very high FDRs, while eve-ttt had a poor statistical power, with TPRs lower and FDRs higher than phyloDE.

### Re-analysis of the empirical dataset from Daunesse *et al*. (2026)

Using the phyloDE method, we performed a re-analysis of the dataset from Daunesse *et al*. (2026). Figure 5 presents the number of differentially expressed genes detected for the two tissues on the three branches from Daunesse *et al*. (2026) for eve-ttt and phyloDE with a Benjamini-Hochberg correction (Benjamini and Hochberg, 1995) of p-values to control for the FDR with a threshold 0.05. The number of DE genes detected by eve-ttt was low to moderate (depending on the design). This result is consistent with our observation on simulated datasets, on which eve-ttt exhibited low statistical power. phyloDE found more DE genes than eve-ttt in all settings, with up to four times as many DE genes for the MR design. To assess whether genes identified as differentially expressed showed consistent changes in their cis-regulatory landscape, we compared their estimated group effects from the RNA-seq analysis to log-fold changes in cis-regulatory element (CRE) activity (Parey *et al*., 2023; Daunesse *et al*., 2026). The number of DE genes with concordant fold-change in the regulatory signal was similar for the two methods, which suggests that phyloDE increases the number of detected DE genes compared to eve-ttt without adding noise. The full lists of DE genes found by phyloDE are reported in the Supplementary Material Section S5.

**Figure 5.**
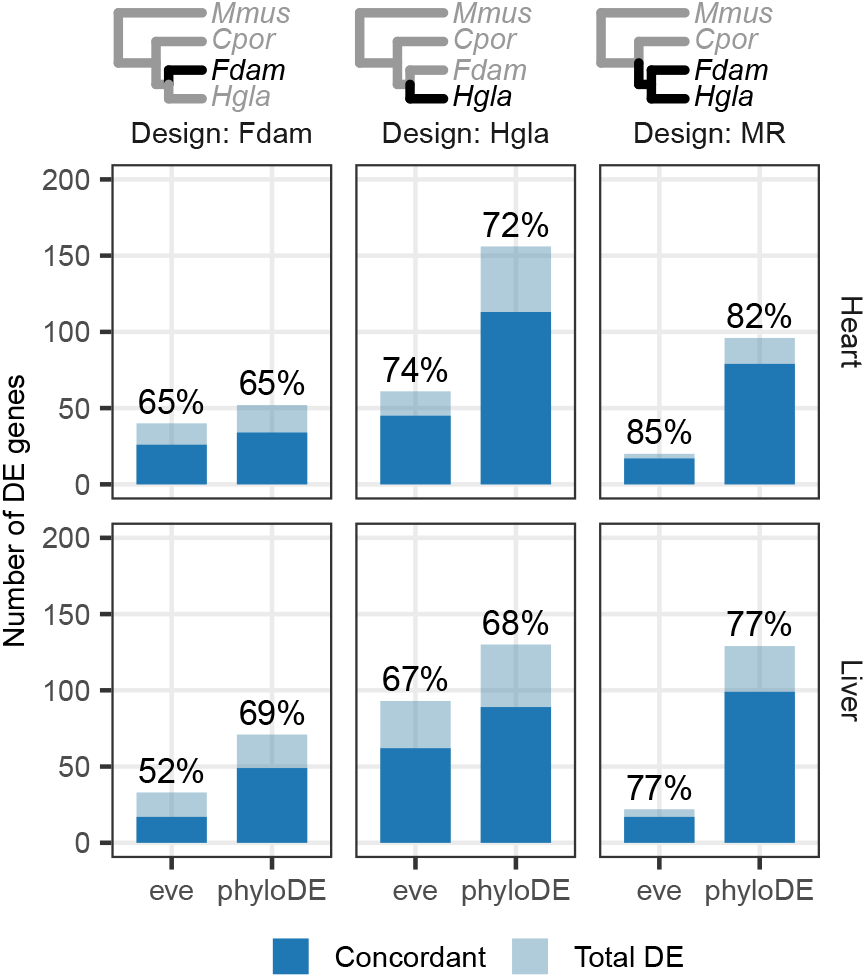
Number of DE genes detected by eve-ttt and phyloDE for differential expression analysis between species *Fdam* versus the rest of the tree (Design: Fdam), between species *Hgla* versus the rest of the tree (Design: Hgla), or between both mole rat species versus the rest of the tree (Design: MR), for the two tissues (Heart, top and Liver, bottom). The two method are eve-ttt (with empirical p-values computed using 20 000 simulations under the null), and phyloDE, with a FDR threshold fixed at 0.05 and Benjamini-Hochberg corrected p-values. The dark blue color indicates the ratio of DE genes that share a concordant signal with the regulatory score. The percentage indicates the proportion of concordant genes among the total number of DE genes.

## Discussion

In this work, we propose phyloDE, a method for phylogenetic differential expression analysis. It is built on three main components: a reasonable trait evolution model, a careful parametrization, and an empirical Bayes regularization that exploits the mean-variance structure of the data. In simulations, we showed that methods that ignore one of the steps above fail in different ways. First, the limma-cor method, that applies regularization but ignores the phylogenetic structure, fails to control the FDR as soon as the data exhibit complex non-independence patterns, in particular when the design of the experiment is confounded with the phylogenetic inertia (in the MR “block” design, see Fig. 4). This result is consistent with Bastide *et al*. (2023). Second, the phylolm method, that uses a relevant OU model but does not regularize the parameters and uses the *t*-tests conditioned on the true values of the parameters being known (see Eq. (2) in the Material and Methods), has a very high level of FDR in our simulation study. This can be explained by the fact that the parameters are indeed poorly estimated, by all methods (see Fig. 1). In this small sample size exemple, the likelihood surface is likely flat, as observed in other studies on the OU (Ho and Ané, 2013; Cornuault, 2022), translating in highly uncertain estimation of the parameters. Any test conditional on these poorly estimated values is therefore bound to be unreliable. The third method, eve-ttt, relies on likelihood ratio tests instead of *t*-tests, and should be more robust to parameter mis-estimation. If the method does control the FDR in some situations, it fails in others (especially for the MR design, see Fig. 2), and, importantly, it can have a statistical power close to zero, detecting a very few number of differentially expressed genes. This low power has already been observed in other studies (Gillard *et al*., 2021; Parey *et al*., 2023; Daunesse *et al*., 2026). Importantly, these flaws, that are particularly pregnant in simulations with a small number of samples such as the ones based on the dataset from Daunesse *et al*. (2026) and showed in the main text (12 samples: 3 replicates on 4 species), do not go away when the number of observations increases. This is demonstrated in the Supplementary Material, on simulations on datasets with the same number of species (4) but more replicates (up to 25 samples: 8 replicates for Hgla, 5 for Fdam, 6 for Cpor, and 6 for Mmus as in the original dataset from Daunesse *et al*. (2026), see Fig. S1); or datasets with more species (up to 17 species, see Fig. S10 and S11). Adding observations, as well as increasing the effect size, tend to improve the TPR for all methods, but does not substancially improve the FDR.

In contrast, the phyloDE method controls the FDR in all the setting we tested, while keeping a reasonable power. It fully benefits from the OU modelling, as using the BM in the same regularized framework led to inflated FDR (see Supplementary Figure S5). It uses a standardized parametrization (see Eq. (5) and (6) in the Material and Methods) with biologically interpretable parameters that lie between 0 and 1, and that lend themselves well to regularization. This strong regularization is justified by the poor individual estimation of these parameters (see Fig. 1 and Supplementary Figure S6), and we showed that the method was robust to a violation of the common process assumption (see Supplementary Figures S2 and S3). phyloDE also benefits from the empirical Bayes regularization techniques specifically developed for RNA-Seq data and implemented in limma (Smyth, 2004; Smyth *et al*., 2005; Phipson *et al*., 2016). In particular, the trend correction, that is similar to the voom correction in a classical setting (Law *et al*., 2014), significantly improves the power of the method on small datasets (see Fig. S4). Finally, batch effects may have a strong effect on gene expression in inter-species studies. Unlike eve-ttt, phyloDE, thanks to its linear model structure similar to limma, can directly include covariables or batch effects in the model itself. In addition, while eve-ttt can only test for lineage-specific shifts, phyloDE can perform differencial analysis for any group design, including paired designs or groups containing several phylogenetically separated clades, such as the “sighted” versus “blind” study of Crayfishes species conducted in Stern and Crandall (2018) (see also the different designs in Bastide *et al*., 2023). We believe these features of the phyloDE method will be particularly usefull for large scale analysis of inter-species gene expression.

On the mole rat dataset, eve-ttt exhibited a low statistical power, consistent with other studies. This low power led researchers to use un-corrected p-values for analyses to recover some signal (Gillard *et al*., 2021; Parey *et al*., 2023; Daunesse *et al*., 2026). In contrast, phyloDE found up to about 150 DE genes on the Heart tissue in the”Hgla” design (Fig. 5), while controlling for the FDR at a 5% threshold using corrected p-values, and retaining consistent biological signal, as exhibited by the percentage of genes with a concordant fold-change in the regulatory signal. Further biological analysis of these lists of genes would be needed to assess the relevance of the selected genes.

Intuitively, the regularization method of phyloDE treats the variance and correlation parameters as nuisance parameters, as the main task of the method is differential expression in the mean. These parameters could however be the focus of other studies, as they can have meaningful biological interpretations of their own. For instance, Stern and Crandall (2018) translate several biological hypotheses on the role of gene expression variation in the evolution of vision loss for crayfishes into hypotheses on the various parameters of the OU. More generally, gene expression data has been used to address numerous other questions such as the construction of species trees based on gene expression, the calculation of organ-specific expression indicators, the inference of ancestral transcriptomic levels, the divergence of expression over time or the study of expression dynamics and evolutionary forces acting on expression (loss through drift, parallel or divergent selection, Brawand *et al*., 2011; Dunn *et al*., 2013; Dimayacyac *et al*., 2023; Gu *et al*., 2019). Of particular interest, the question of detecting of shifts on the evolutionary rate of the process, instead of shifts in the mean, has been tackled, e.g., by the betaSharedTest method of EVE (Rohlfs and Nielsen, 2015), or the CAGEE method Bertram *et al*. (2022). In a non-phylogenetic RNA-Seq context, such tests on the variance have been explored, relying, e.g., on the Levene test (Phipson and Oshlack, 2014). Adapting them to the inter-species framework could be an interesting future research avenue.

The phyloDE method to detect shifts in the mean could be improved in several ways. First, just as the approach of Smyth *et al*. (2005), it relies on a two steps procedure, first regularizing the structural variance parameters *λ* and *ρ*, and then applying an empirical Bayes procedure to the de-correlated data. Although this approach is computationally efficient and works well when the dataset contains many genes, an integrated, one-step approach might be desirable to avoid any error propagation. The phylogenetic empirical Bayes procedure of Montoya *et al*. (2026) for high dimensional datasets might be a promising step in this direction. Second, because it relies on a transformation of the tree to fit the OU process just like phylolm (see Material and Methods), phyloDE can only handle trees that are ultrametric, such as trees dated in calendar years and containing extent species only. Third, phyloDE, just as limma, relies on a pre-processing normalization step of the count data. In classical RNA-Seq studies, methods based on the direct modelling of count data such as edgeR (Robinson *et al*., 2009) or DESeq2 (Love *et al*., 2014) have been highly successful. They rely on a variance-inflated negative binomial distribution model, that we showed could be extended in a phylogenetic context with a multivariate Poisson log-normal (PLN) model (Bastide *et al*., 2023), the distribution used in compcodeR for phylogenetic simulations. Beyond simulations, this model could be used for direct data analysis, drawing from the now well established software for PLN data fitting in various context (Gallopin *et al*., 2013; Zhang *et al*., 2015; Chiquet *et al*., 2019, 2021), including zero-inflated versions (Batardière *et al*., 2025). Finally, we focussed in this work on RNA-seq datasets, but the model could be used to analyze other types of omic datasets. In Daunesse *et al*. (2026), the authors explore the joint analyses of gene expression and gene regulation. Developing a multi-omics approaches combining several omic layers across species is another promising perspective.

## Materials and methods

### The phyloDE method

The phyloDE method relies on a phylogenetic linear regression with moderated *t*-statistics and regularized parameters, as detailed below.

#### Setting, notations and normalization

We assume that we have access to *n*_*i*_ individual measurements for each of the 1 *≤ i ≤ n* species of the studies, with a total 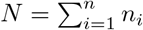 number of samples. RNA-Seq data are typically discrete counts, but we assume here that it has been properly normalized, so that it can be treated as a continuous measurement. In the following, we use as a default the log_2_ transformation of the transcripts per million (TPM, Wagner *et al*., 2012) scores, with normalization factors computed with the Trimmed Mean of M-values from Robinson and Oshlack (2010). This transformation was shown to behave well in a phylogenetic context (Bastide *et al*., 2023), but any other normalisation method taking the species-specific gene lengths *l*_*gi*_ into account could be employed, depending on the specificities of the data at hand. We write *Y*_*gi*_ the RNA-Seq normalized data for gene 1 *≤ g ≤ p* and sample 1 *≤ i ≤ N*. Our goal is to test for differential gene expression between two groups of species, or more generally to test the coefficients associated with a model (or design) matrix **X** (with *N* lines and *q* columns) in a linear model.

#### Phylogenetic regression

In a phylogenetic regression, we model each vector **Y**_*g*_ of the *N* normalized measurements for gene *g* with a linear model:

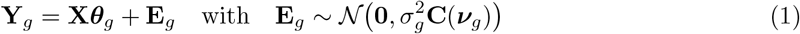

where ***θ***_*g*_ a vector of *q* coefficients, and **E**_*g*_ is a centered Gaussian random vector with total variance 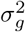 and structured correlation matrix **C**(***v***_*g*_), that depends on a vector of parameters ***v***_*g*_, and that represents the correlation both between species and between samples inside a species. Testing for differential expression then amounts to testing for the nullity of coefficient *θ*_*g,k*_ associated with the grouping column **X**_*k*_ of the design matrix. If we assume that the parameters ***v***_*g*_ are known, then the *t*-statistic

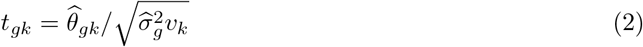

follows a Student *t* distribution with *d*_*g*_ = *N − q* degrees of freedom (when there is no missing data). The statistic relies on the generalized least square estimate of the coefficient 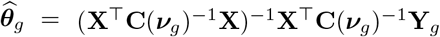 and the restricted maximum likelihood (REML, see, e.g., Searle *et al*. 1992) of the variance 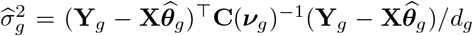, with *v*_*k*_ is the *k*^*th*^ diagonal entry of (**X**^⊤^**C**(***v***_*g*_)^*−*1^**X**)^*−*1^. By default, this is the statistic that is used in phylolm (Ho and Ané, 2014a), that implements a linear time algorithm to compute it (avoiding the computational cost of inverting **C**(***v***_*g*_)).

#### Challenges with gene expression data

As mentioned in the Result Section, we face four main challenges when trying to use the phylogenetic framework and the *t*-statistics above in a differential gene expression analysis using RNA-Seq data, that are recalled below. (i) Matrix **C**(***v***_*g*_) must represents the correlations induced by a biologically meaningful process of gene evolution. (ii) The parameters ***v***_*g*_ are actually unknown, and we know that they were poorly estimated (see Fig. 1 and Supplementary Material Section S3), so that they must be carefully chosen and regularized. (iii) We usually have access to several thousands of genes, so this analysis must be repeated many times, and the tests need to be moderated through a regularization of the variance that shares information between genes to gain some statistical power. (iv) RNA-Seq data has some technical specificities, including a possible mean-variance relationship. We propose solutions to these four challenges in the paragraphs below.

##### (i) Evolution process: the OU with within-species variation

Phylogenetic comparative methods use stochastic models of trait evolution such as the Brownian Motion (BM; Felsenstein, 1985) or the Ornstein-Uhlenbeck (OU; Hansen and Martins, 1996) to derive correlations between species (for reviews, see, e.g., Felsenstein 2004; Harmon 2019). These methods typically assume that the tree is fixed and known, and we further assume that it is ultrametric of total height *t*. The Ornstein-Uhlenbeck model has been extensively used in a gene expression context (Roth *et al*., 2014; Rohlfs and Nielsen, 2015; Dimayacyac *et al*., 2023). For each gene *g*, in addition to a rate evolution variance 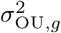 and a within-species variance 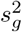, it has an extra parameter *α*_*g*_, that controls the strength of the pull towards the optimal trait value. Assuming a fixed root, the covariance between samples 1 *≤ i, j ≤ N* can be written as (see, e.g., Ho and Ané, 2014b):

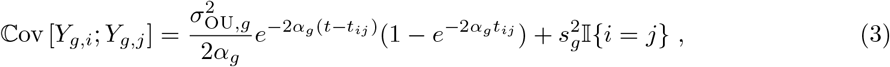

where *t*_*ij*_ the time between the root of the tree and the most recent common ancestor of *i* and *j*, and I*{i* = *j}* = 1 if *i* = *j*, and 0 otherwise. In this traditional formula (3) for the covariance matrix, the natural parameters 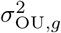, *α*_*g*_ and 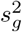 impact both the total amount of variance and the correlation structure between individuals, making it difficult to grasp the concrete effect of one parameter varying. To alleviate this issue, we cast this covariance structure into the linear model framework of Eq. (1), that clearly separates the total variance parameter 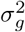 from the correlation structure **C**(***v***_*g*_). Using the usual re-scaled time 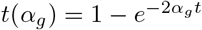 (Blomberg *et al*., 2003; Ho and Ané, 2014a), we write:

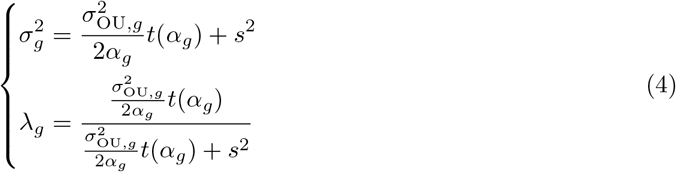

so that 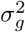 is the total variance, and the structure matrix depends on the two parameters (*λ*_*g*_, *α*_*g*_) as:

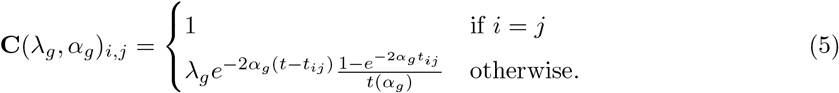

This parametrization has several interesting properties. First, the parameter *λ*_*g*_ directly captures the within-species correlations: if *i* and *j* are two samples from a single species, then *t*_*ij*_ = *t*, and **C**(*λ*_*g*_, *α*_*g*_)_*i,j*_ = *λ*_*g*_. Second, we recover two well-known models as limit cases when *α*_*g*_ varies:

- when *α*_*g*_ *→* 0, then for *i≠ j*, **C**(*λ*_*g*_, 0)_*i,j*_ = *λ*_*g*_*t*_*ij*_*/t* is the correlation structure of a BM;
- when *α*_*g*_ *→* +*∞*, **C**(*λ*_*g*_, *α*_*g*_)_*i,j*_ *→* 0 if *i j* are from *different* species, and **C**(*λ*_*g*_, *α*_*g*_)_*i,j*_ *→ λ*_*g*_ if *i≠ j* are from *the same* species: the correlation matrix has a species-block structure, that can be captured using the limma-cor method.

In this framework, the OU model can hence be seen as a natural extension of limma-cor, but with a different correlation structure. These limma-cor and BM models are described in more details in Supplementary Material Section S1.

##### (ii) Parametrization: the OU normalized correlation structure

While parameter *λ*_*g*_ has a clear interpretation in term of correlation structure, the effect of *α*_*g*_ on the correlation is less obvious. Further, *α*_*g*_ is notoriously difficult to estimate, even for large trees (Ho and Ané, 2013). The likelihood surface tends to exhibit some “ridge” regions in 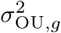 and *α*_*g*_ (Cornuault, 2022). To alleviate theses issues, Cornuault (2022) proposes to use the *ρ* metric, defined as:

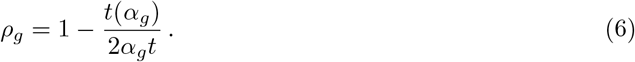

It can be interpreted as representing the percent decrease in trait variance caused by the OU as compared to the variance expected under under BM (Cornuault, 2022), and has natural variations between 0 (*α*_*g*_ *→* 0, BM-like) and 1 (*α*_*g*_ *→* +*∞*, no tree structure). As the function defining *ρ*_*g*_ is strictly increasing for *α*_*g*_ *≥* 0, it can be inverted, so that the structure matrix **C** can be written as a function of *λ*_*g*_ and *ρ*_*g*_ only (see Supplementary Material Equation (S5)). This is the base parametrization used in phyloDE. Although it appears to be less natural than the usual covariance matrix expression (3), it formulates the OU induced structure as a normalized correlation matrix, with ones on the diagonal, and parameters *λ*_*g*_ and *ρ*_*g*_ that vary between 0 and 1. If the tree is ultrametric, these parameters can also be seen as defining a branch length transformation of the original tree similar to Pagel’s *λ* (Pagel, 1999), clarifying their interpretation in term of within-species and between species correlations (see Fig. 3 and Supplementary Material Equation S6). Importantly, all transformed trees have a unit height, so that they actually capture the correlation structure, while the total scaling variance is captured by parameter 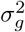 from Eq. (4). In the practical implementation, we numerically inverse (6) to solve for *α*_*g*_, and then apply standard tree transformation functions from phylolm (Ho and Ané, 2014a) and re-scale it to unit height to compute the correlation matrix.

##### (ii-bis) Regularization of the OU structure parameters *λ*_*g*_ and *ρ*_*g*_

Crucially, this parametrization lends itself better to regularization, as the parameters lie on a common scale for all the genes. Using a technique similar to the limma-cor method (Smyth *et al*., 2005), we make the assumption that all the genes share the same parameters *λ*_0_ and *ρ*_0_. As in Smyth *et al*. (2005), assuming that we have estimated 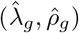 independently for each gene *g* (using, e.g., the REML estimates of phylolm), phyloDE then robustly estimates *λ*_0_ and *ρ*_0_ by the trimmed mean of all these estimates in the inverse hyperbolic tangent transformed space: 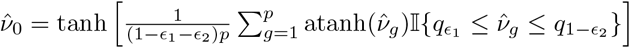, where ***v*** represents either *λ* or *ρ*, and 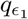 and 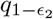 are the quantiles of the sample 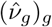. In Smyth *et al*. (2005), default trimming proportions are *ϵ*_1_ = *ϵ*_2_ = 0.15. Here, as we are primarily concerned with controlling the FDR, we set asymmetric proportions *ϵ*_1_ = 0.25 and *ϵ*_2_ = 0.05, favoring a stronger phylogenetic structure that tend to discard lower shifts as random drift (see Supplementary Figure S9). phyloDE then follows the procedure of Smyth *et al*. (2005), that treats the regularized estimated correlation parameter as known and uniform across genes, and computes the *t*-statistics (2) conditionally on this known values (by injecting 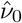 instead of ***v***_*g*_ in the associated estimates). In a non-phylogenetic context, Smyth *et al*. (2005) show that the procedure works well for microarray experiments with within-arrays replicate spots, and that the truncated mean, although less efficient than a full maximum likelihood approach, is sufficient when the number of genes is large. They also demonstrate that the strong regularization on the correlation parameter is needed to gain some statistical power when combined to the empirical Bayes regularization of the variance explained below.

##### (iii) Moderated *t*-statistics: empirical Bayes regularization of the variance

Assuming that the correlation structure 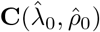 is known, the *t*-statistics (2) still depends on the gene-specific estimated variance parameters 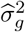. phyloDE then employs the empirical Bayes procedure implemented in the popular R package limma (Ritchie *et al*., 2015), and performs a further regularization of the variance estimate across the genes. It starts by assuming a common prior conjugate inverse gamma distribution 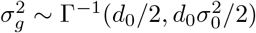 on all the variance parameters, so that the posterior expectation of 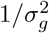 given 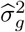 is 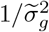, with 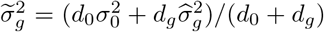. Smyth (2004) then shows that the moderated *t*-statistic 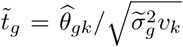 follows a centered *t* distribution with *d*_0_ + *d*_*g*_ under the null hypothesis *θ*_*gk*_ = 0. If 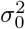 and *d*_0_ are known, using the moderated statistics generally improves the power of the test, increasing the degrees of freedom from *d*_*g*_ to *d*_0_ + *d*_*g*_. These hyper-parameters 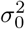 and *d*_0_ are unknown, but can be robustly estimated from the pool of the estimated variances 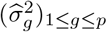 on all the genes, by noting that, under the hierarchical model, each estimated variance is marginally distributed as a scaled Fisher distribution 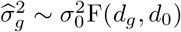 (Phipson *et al*., 2016).

##### (iv) RNA-Seq data: leveraging the mean-variance relationship

In RNA-Seq experiment, it is common to observe a mean-variance relationship, with highly expressed genes tending to have a lower variance. Law *et al*. (2014) tackle this characteristic either using pre-processed weights (in the voom methodology), or by adding a trend in the empirical Bayes procedure (limma-trend), and show that both perform similarly. The limma-trend method replaces the common prior on the variances presented above by a gene-specific prior 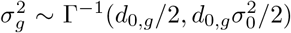, with *d*_0,*g*_ a smooth function of *A*_*g*_ the mean normalized expression of gene *g*. The log-variances 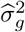 can then be detrended using a (robust) lowess regression against *A*_*g*_ (Phipson *et al*., 2016), before applying the empirical Bayes method presented above. This trend correction can be readily applied in a phylogenetic regression context, and we showed that it indeed improved the power of our phyloDE method.

#### Comparison with the EVE method

The Expression Variance and Evolution (EVE) model (Rohlfs *et al*., 2014; Rohlfs and Nielsen, 2015) is similar to phyloDE in that it uses a OU model on the tree to account for phylogenetic correlations to perform a phylogenetic ANOVA in the context of gene expression analysis, but it differs in several crucial ways, detailed below: parametrization, regularization, and testing procedure. On the variance structure, the main difference is that they set 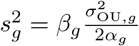 in Eq. (3), and estimate parameters *β*_*g*_, 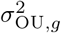 and *α*_*g*_ for each gene independently, without regularization. For the group structure, instead of relying on the linear model of Eq. (1), they directly model the expectation of the trait by assuming that there are two groups, and that each group has its own optimal value parameter *θ*_*k*_. As only the expectation at the tips of the tree is identifiable for a OU on an ultrametric tree (Ho and Ané, 2014b), it is equivalent to a linear model with a design matrix **X** coding for the group structure. Note that this approach is less flexible than the phyloDE method based on the linear model, as it cannot directly include covariables or batch effects. The test for differential expression is then carried using a likelihood ratio test, comparing a model with *θ*_1_ = *θ*_2_ versus a model with *θ*_1_*≠ θ*_2_. The p-values are obtained from the empirical distribution of the test statistic under the null, obtained through simulations.

### Simulation studies

#### Tree and design

In all simulations, we used the empirical mole rats tree, that has *n* = 4 species: the naked mole-rat (*Heterocephalus glaber*, Hgla), the Damaraland mole-rat (*Fukomys damarensis*, Fdam), the guinea pig (*Cavia porcellus*, Cpor), and the mouse (*Mus musculus*, Mmus). We extracted this dated tree from the larger mammals tree from Fritz *et al*. (2009), as curated in Hassler *et al*. (2020). We assumed three replicate measurements per species, so that the total number of samples was *N* = 12. We used two different designs, both defining groups of species (see Fig. 4, top row). In the “Hgla” design, the replicates of the naked mole-rat (Hgla) where in one group, while all other replicates from other species where in the other. In the “MR” design, the measurements from both species of mole rats (Hgla and Fdam) were in one group, while the replicates from the two other species (Cpor and Mmus) were in the other. Note that, while both designs are structured in “blocks”, i.e., are consistent with clades on the tree, the first design separates the two mole rats species, that are closely related on the tree, in two different groups.

#### Gene expression dataset

The dataset used for empirical simulations below was extracted from Daunesse *et al*. (2026). We used the raw counts matrix for liver expression provided by the authors upon request. Gene length extraction was performed using the Ensembl BioMart web tool on October, 2025 for the guinea pig, the mouse and the naked mole-rate (Dyer *et al*., 2025). We used the naked mole-rate gene lengths as a proxy for the Damaraland mole-rate gene lengths, given that the two species are close. Note that this data was used to set the numerical values of the parameters in our simulation studies, not for the real dataset analysis. In all the simulations presented in the main text, we kept only three replicates per species. Simulations using the full number of replicates per species from Daunesse *et al*. (2026) (i.e., 8 replicates for Hgla, 5 for Fdam, 6 for Cpor, and 6 for Mmus) are presented in the Supplementary Material.

#### Gaussian simulations

We simulated normalized datasets according to the Gaussian linear model (1), with a OU structure matrix as in (5), with total variance fixed to 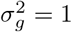 in all settings, on the mole rat tree rescaled to unit height. We simulated *p* = 10 000 genes independently, with fixed parameters *λ* and *α* across all the genes, and varied *λ* from 0.1 (low phylogenetic signal, samples inside a species are almost independent) to 0.9 (strong phylogenetic signal, samples inside a species are strongly correlated), and *α* from 0.14 (*ρ* = 0.13, species are strongly correlated as in a BM), to 13.9 (*ρ* = 0.96, species are almost independent). When varying *λ, α* was fixed to 1.39 (*ρ* = 0.66), while when varying *α, λ* was fixed to 0.75. The design matrix **X** in linear model (1) reflected the two “Hgla” and “MR” designs, and we took a fixed effect size of 4, with 150 truly differentially expressed genes. Each setting was replicated 20 times.

#### Empirical count simulations

We used the R package compcodeR (Soneson, 2014) to simulate realistic count data as in Bastide *et al*. (2023), taking the liver mole rats dataset with the “MR” design as a base, filtering out genes with low counts (less than 100 counts across all measurements), which left us with 8 455 genes. The simulation procedure is described in details in Bastide *et al*. (2023). The package simulates count data *C*_*gi*_ using a Poisson log-normal distribution:

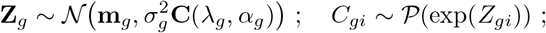

were **C**(*λ*_*g*_, *α*_*g*_) is as in (5), and parameters 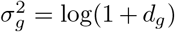 and *m*_*gi*_ = log(*µ*_*gi*_) *−* 0.5 *·* log(1 + *d*_*g*_) are chosen so that the expectation and variance of the *C*_*gi*_ match those of a Negative Binomial distribution with mean *µ*_*gi*_ and dispersion *d*_*g*_, further assuming that the mean is structured as 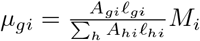, with *A*_*gi*_ the true expression level of gene *g* in sample *i, l*_*gi*_ the species specific gene length and *M*_*i*_ the sampling depth for sample *i* (Robinson and Oshlack, 2010). Differential expression between two groups is achieved by constraining *A*_*gi*_ to be either equal to *A*_*g*1_ if *i* is in group 1 or *A*_*g*2_ if *i* is in group 2, with 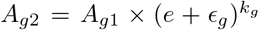, with *e* the minimum effect size, *ϵ*_*g*_ random variables independent identically distributed according to an exponential distribution with parameter 1, and *k*_*g*_ = 0 if the gene *g* is not differentially expressed, *k*_*g*_ = 1 if it is up-regulated, and *k*_*g*_ = *−*1 if it is down-regulated. The base expression level *A*_*g*,1_ and the dispersion *d*_*g*_ were empirically estimated from the dataset, and the sequencing depth *M*_*i*_ was randomly drawn for each sample from an uniform distribution with bounds the observed empirical minimal and maximal values of the library size across all samples. To take the phylogenetic structure into account, we estimated global values for 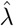 and 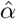 by taking the median of estimates on all genes using phylolm with the OU model on log_2_TPM normalized data. We then tested three simulation settings. In the “fixed parameter” setting, all the genes were simulated using the same parameters 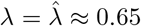 and 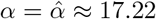 This is the most favorable case for our method, as it regularizes these estimates to a single value. Note that *α* was quite hight (*ρ* = 0.97), so that the species were only weakly correlated in this base simulation scenario. In the “low variance” setting, we generated a different *λ*_*g*_ and *α*_*g*_, all drawn from, respectively, a beta distribution with mean 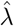 and standard error 0.05 and a gamma distribution with mean 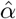 and standard error 1. Finally, we did the same in the “high variance” setting, but with standards errors, respectively, of 0.1 and 5 (see Supplementary Figure S2). In the high variance case, this lead to simulation parameters *λ*_*g*_ varying between 0.22 and 0.93, and *α*_*g*_ between 4.83 and 40.05, covering a relatively high range of parameters. In all cases, we used a base effect size of *e* = 4, with 150 differentially expressed genes out of the 8 455 simulated genes. Each setting was replicated 20 times.

#### Methods used for differential expression analysis

We compared four main methods for differential expression analysis: limma-cor, eve-ttt, phylolm and phyloDE. We did not include edgeR or DESeq2, as previous study showed that these methods were not robust to phylogenetic noise (Bastide *et al*., 2023). For limma-cor, we used version 3.66 of package limma available on bioconductor, with empirical Bayes moderation. For eve-ttt, we used the R implementation described in Gillard *et al*. (2021), and available on gitlab (https://gitlab.com/sandve-lab/evemodel). As advocated in the tutorial and in recent studies using this method (Parey *et al*., 2023; Daunesse *et al*., 2026), p-values were obtained from the empirical distribution of the test statistic under 20 000 simulations of the null hypothesis. For phylolm, we used the “OUfixedRoot” model with added measurement error on the tree with individual replicates as in Bastide *et al*. (2023). Finally, we used phyloDE as presented above with the OU model, regularization of parameters, and empirical Bayes moderation of *t*-values. For limma-cor and phyloDE, we used the trend correction in the empirical count data simulations only. All p-values were corrected using the Benjamini-Hochberg procedure (Benjamini and Hochberg, 1995) with R function p.adjust (R Core Team, 2025). Normalized inference scripts were included into the compcodeR package (branch phyloDE, https://github.com/pbastide/compcodeR/tree/phyloDE). Trees were handled with packages ape (Paradis *et al*., 2004) and phytools (Revell, 2012), and plots with package ggplot2 (Wickham, 2009) and cowplot (Wilke, 2025).

### Re-analysis of the empirical dataset from Daunesse ***et al***. (2026)

#### Setting

To perform a re-analysis of the liver gene expression, we used the count data already normalized and log-2 transformed with the effective transcript length strategy described in Daunesse *et al*. (2026), and available online at https://gitlab.com/maelle.daunesse/daunesse_et_al_2025. We used the same species tree as in the simulation study, and kept all the replicates per species from Daunesse *et al*. (2026) (from 5 to 8, see above). We analyzed both the liver and heart data, with the three possible designs: “MR”, “Hgla” and “Fdam”, and using phyloDE and eve-ttt with the same parameters as in the simulation studies.

#### Concordance with cis-regulatory element activity

To assess whether genes identified as differentially expressed show consistent changes in their cis-regulatory landscape, we compared their estimated group effects from the RNA-seq analysis to log-fold changes in cis-regulatory element (CRE) activity. CRE activity scores were taken from Daunesse *et al*. (2026), where they were computed for each gene as the quantile-normalised sum of H3K27ac ChIP-seq activity across promoters (*±*10 kb of the TSS) and enhancers (*±*100 kb of the TSS), in heart and liver tissues, across the four species. For each gene, tissue and branch contrast (Hgla and MR), we computed the CRE log-fold change as 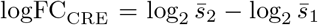, where 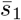 and 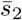 denote the mean CRE activity scores across species in groups 1 and 2, respectively (with a small offset of 0.01 added prior to the log transform to avoid undefined values). On the RNA-seq side, we used the estimated group effect 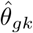 from phyloDE (or the equivalent 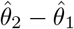 from eve-ttt), which captures the inferred difference in expression optima between groups under the phylogenetic model. For each method, we restricted the analysis to genes called as differentially expressed and checked whether the sign of 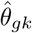 was concordant with the sign of logFC_CRE_.

## Supplementary Material

Supplementary Material contains some precisions about the methods (S1); supplementary simulation results (S2); precisions about the estimation example on one dataset (S3); a study of the effects of some hyper-parameters of the methods (S4); the full list of DE genes found by phyloDE in the empirical anlysis (S4); and supplementary results from the reanalysis of the empirical dataset (S5).

## Acknowledgments

We are grateful to the INRAE MIGALE bioinformatics facility (MIGALE, INRAE, 2020. Migale bioinformatics Facility, doi: 10.15454/1.5572390655343293E12) for providing computing and storage resources. AL received funding from the European Union (ERC, PROMISE, 101087830). Views and opinions expressed are however those of the author(s) only and do not necessarily reflect those of the European Union or the European Research Council. Neither the European Union nor the granting authority can be held responsible for them. For the purpose of open access, a CC-BY public copyright licence has been applied by the authors to the present document and will be applied to all subsequent versions up to the Author Accepted Manuscript arising from this submission.

## Data Availability

The R package phyloDE is open-source and available at https://github.com/pbastide/phyloDE, with documentation website: https://pbastide.github.io/phyloDE/. All the scripts and data to reproduce the analyses of the paper are available at https://github.com/i2bc/phyloDE_paper2026.

## S1 Supplementary methods

In the main text, we presented the phyloDE method with the OU process for differential expression analysis. In this Supplementary Matterial Section, we give some further details on the limma-cor method and on the phyloDE method using the BM. We clarify that the three methods can be seen as an implementation of the same framework, but using with different structures for the covariance matrix.

### Setting and notations

We assume that we have access to *n*_*i*_ individual measurements for each of the 1 *≤ i ≤ n* species of the studies, with a total 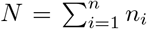 number of samples. We write *C*_*gi*_ the RNA-Seq count data for gene 1 *≤ g ≤ p* and sample 1 *≤ i ≤ N*. We also assume that we have access to sample specific normalisation factors *m*_*i*_ (computed, e.g., through Relative Log Expression, Anders and Huber 2010 or Trimmed Mean of M-values, Robinson and Oshlack 2010, see Dillies *et al*. 2013 for a review), as well as gene lengths *l*_*gi*_ for each gene *g* in sample *i*, that might vary across species. Our goal is to test for differential gene expression between two groups of species, or more generally to test the coefficients associated with a model (or design) matrix **X** (with *N* lines and *q* columns) in a (generalized) linear model. We denote by *Y*_*gi*_ the normalized log-transformed data for gene 1 *≤ g ≤ p* and sample 1 *≤ i ≤ N*.

### The limma-cor method: limma with within-species correlations

In a non-phylogenetic context, the popular R package limma (Ritchie *et al*., 2015) models each vector **Y**_*g*_ of the *N* normalized measurements for gene *g* with a linear model: **Y**_*g*_ = **X*θ***_*g*_ + **E**_*g*_ with 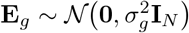 where **E**_*g*_ is a vector of independent identically distributed (iid) centered Gaussian random variables with variance 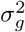, and ***θ***_*g*_ a vector of *q* coefficients associated with the design matrix **X**. The limma method uses an empirical Bayes approach to regularize the variance parameters 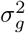, and computes moderated t-test statistics (Smyth, 2004) combined with a global multiple testing correction (Benjamini and Hochberg, 1995) to perform differential expression analysis between groups of samples for each gene. To adapt this method to the phylogenetic context, it is reasonable to assume that the individual measurements within a single species are correlated, even if species in themselves are assumed to be independent. This leads to assuming a linear model with structured errors:

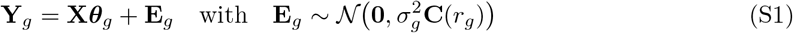

where **C**(*r*_*g*_) is a bloc-matrix correlation structure that is such that:

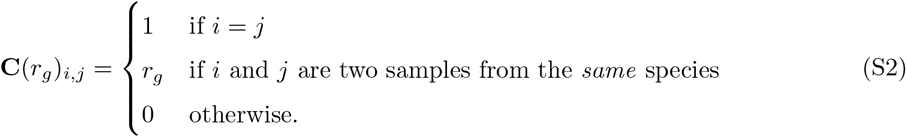

Smyth *et al*. (2005) proposes an extension of the limma method in this setting in a three-steps procedure: (i) estimate the parameters of the linear model 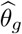, 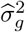 and 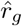 independently for each gene (using a mixed model REML approach); (ii) regularize the correlation coefficients by assuming that all the genes have the same parameter *r*, and robustly estimating 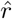 by the trimmed mean of all the 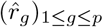 in the inverse hyperbolic tangent transformed space, as implemented in the duplicateCorrelation function from the R limma package; and (iii) apply the original empirical Bayes procedure to the decorrelated data

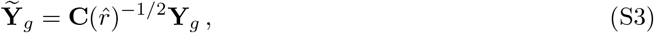

with 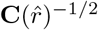 such that 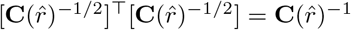. Smyth *et al*. (2005) show that the procedure works well for microarray experiments with within-arrays replicate spots. They also demonstrate that the strong regularization on the correlation parameter *r*_*g*_ is needed to gain some statistical power when combined to the empirical Bayes regularization of the variance, and that the truncated mean of step (ii), although less efficient than a full maximum likelihood approach, is sufficient when the number of genes is large. We use the same approach in the trimmed mean of the main text. We refer to this method as limma-cor.

### The phyloDE BM method: using a BM with within-species variation instead of a OU process

For any gene 1 *≤ g ≤ p*, the BM with evolutionary rate 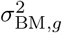 on the ultrametric tree with additional within-species variance 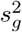 can be cast into the linear regression framework of Eq. (1) from the main text, with a structure matrix **C**(*λ*_*g*_) defined as:

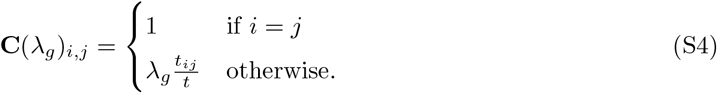

where, for any two samples 1 *≤ i, j ≤ N*, *t*_*ij*_ is the time between the root of the tree and the most recent common ancestor of *i* and *j*. As in the main text, we assume that the tree is fixed and known, and that it is ultrametric with total height *t*, so that for any two samples *i* and *j* that are in the same species, *t*_*ij*_ = *t*. The total variance of the linear model of Eq. (1) from the main text is then equal to 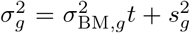, and the parameter is 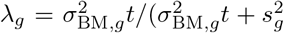, which is equivalent to Pagel’s *λ* parameter (Pagel, 1999, see, e.g., Leventhal and Bonhoeffer, 2016). Note that, if *i* and *j* are two samples from the same species, **C**(*λ*_*g*_)_*i,j*_ = *λ*_*g*_, so that *λ*_*g*_ can be seen as a correlation coefficient similar to *r*_*g*_ defined in equation (S2). Further, when the tree is a star-tree, then *t*_*ij*_ is 0 if *i* and *j* are not in the same species, and *t* otherwise, so that the structure matrix becomes equivalent to (S2), except that *λ*_*g*_ is constrained to be between 0 and 1. Once the parameter *λ*_*g*_ is estimated for each genes, we follow the same procedure as limma-cor: we first regularize the parameters into a global estimate 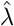 through the trimmed mean as described in the main text; and then apply the standard limma empirical Bayes procedure on the de-correlated data, using 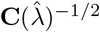.

#### phyloDE OU: closed form formulas for the *ρ* parametrization

Using W_0_ the principal Lambert W function (Cornuault, 2022), the correlation matrix of Eq. (5) of the main text can be formally written as a function of *λ*_*g*_ and *ρ*_*g*_ only:

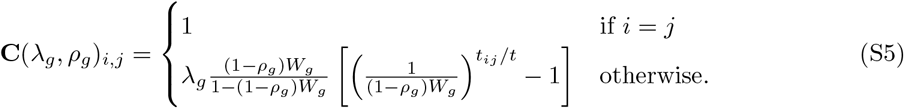

Where 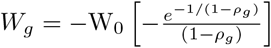. A OU with parameters *λ*_*g*_ and *ρ*_*g*_ on an ultrametric tree with height *t* induces the same correlation structure as a BM on a tree with branch lengths modified as:

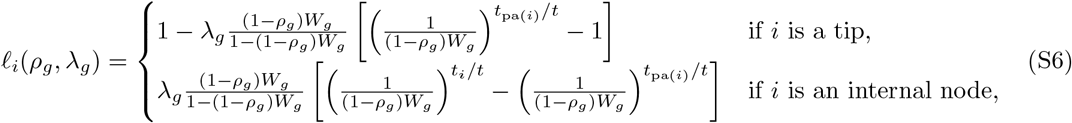

where *l*_*i*_(*ρ*_*g*_, *λ*_*g*_) is the length of the branch ending at node *i* in the modified tree, and *t*_*i*_ and *t*_pa(*i*)_ are the times between the root and, respectivelly, node *i* and its parent node pa(*i*) in the original tree.

## S2 Supplementary simulation results

**Figure S1:**
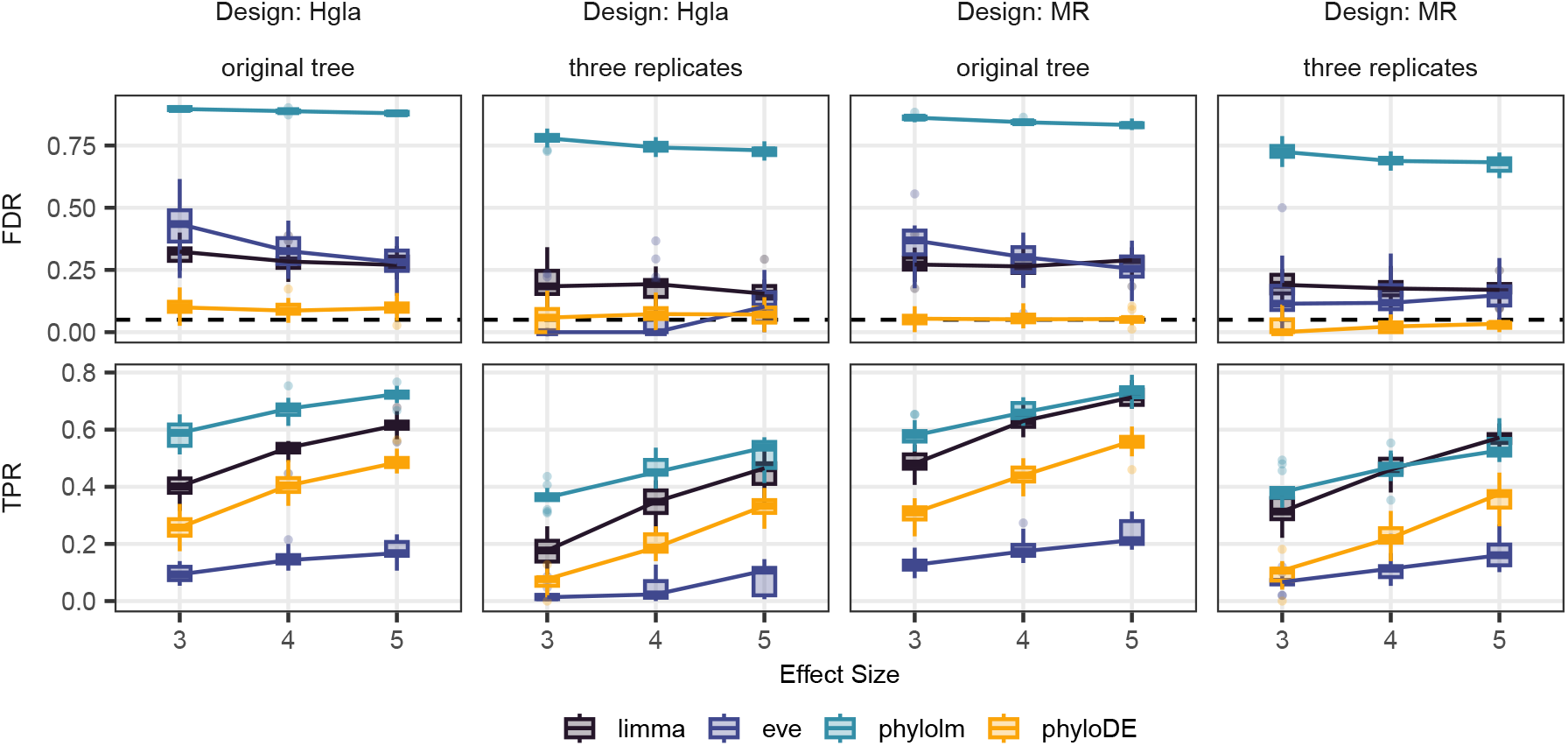
False Discovery Rate (FDR, top row) and True Positive Rate (TPR, bottom row) using Benjamini-Hochberg corrected p-values with a threshold of 5% (dashed horizontal line) for differential expression analysis between species *Hgla* versus the rest of the tree (Design: Hgla, left two columns), or both mole rat species versus the rest of the tree (Design: MR, right two columns), for 20 datasets simulated with the OU process using compcodeR on the empirical tree either with the original number of replicates (left) or with 3 replicates only per species (right), and using empirical parameters for all genes (“fixed”), with an effect size varying from 3 to 5. Methods are limma (with block correlation and empirical Bayes correction of fitted variances with trend), eve-ttt (with empirical p-values computed using 20 000 simulations under the null), phylolm (assuming a OU process with a fixed root), and phyloDE (assuming a OU process with a fixed root and empirical Bayes correction of fitted variances with trend). The design with only three replicates is more difficult for all methods, and increased effect size leads to better TPR.

**Figure S2:**
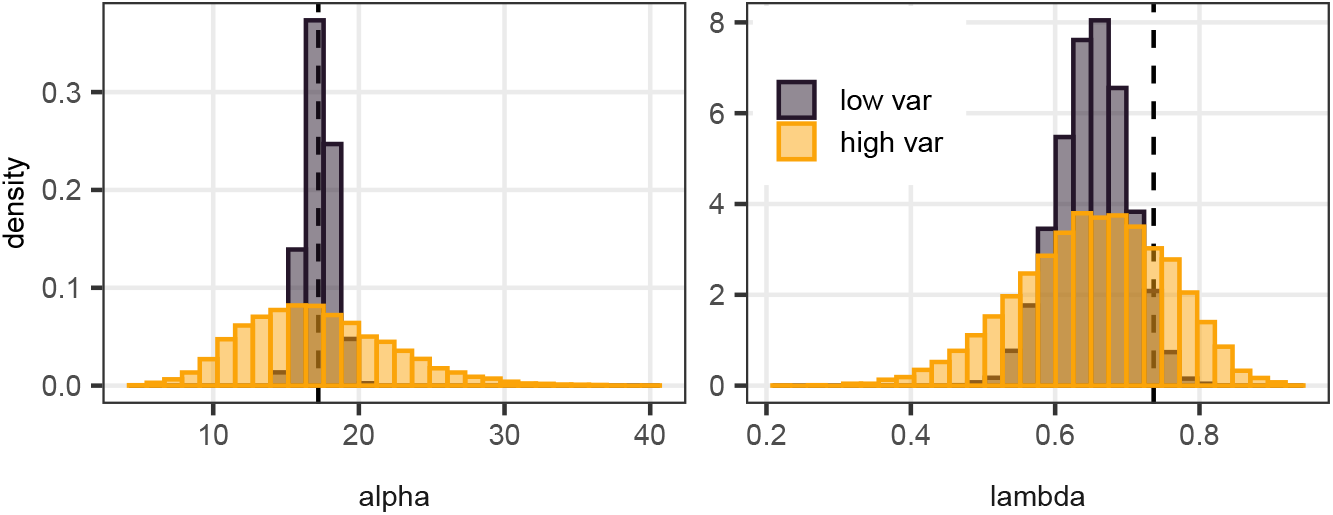
Distribution of the parameters over the genes used in the “fixed” (dashed line), “low var” (dark grey) and “high var” (light orange) settings for parameters *α*_*g*_ and *λ*_*g*_.

**Figure S3:**
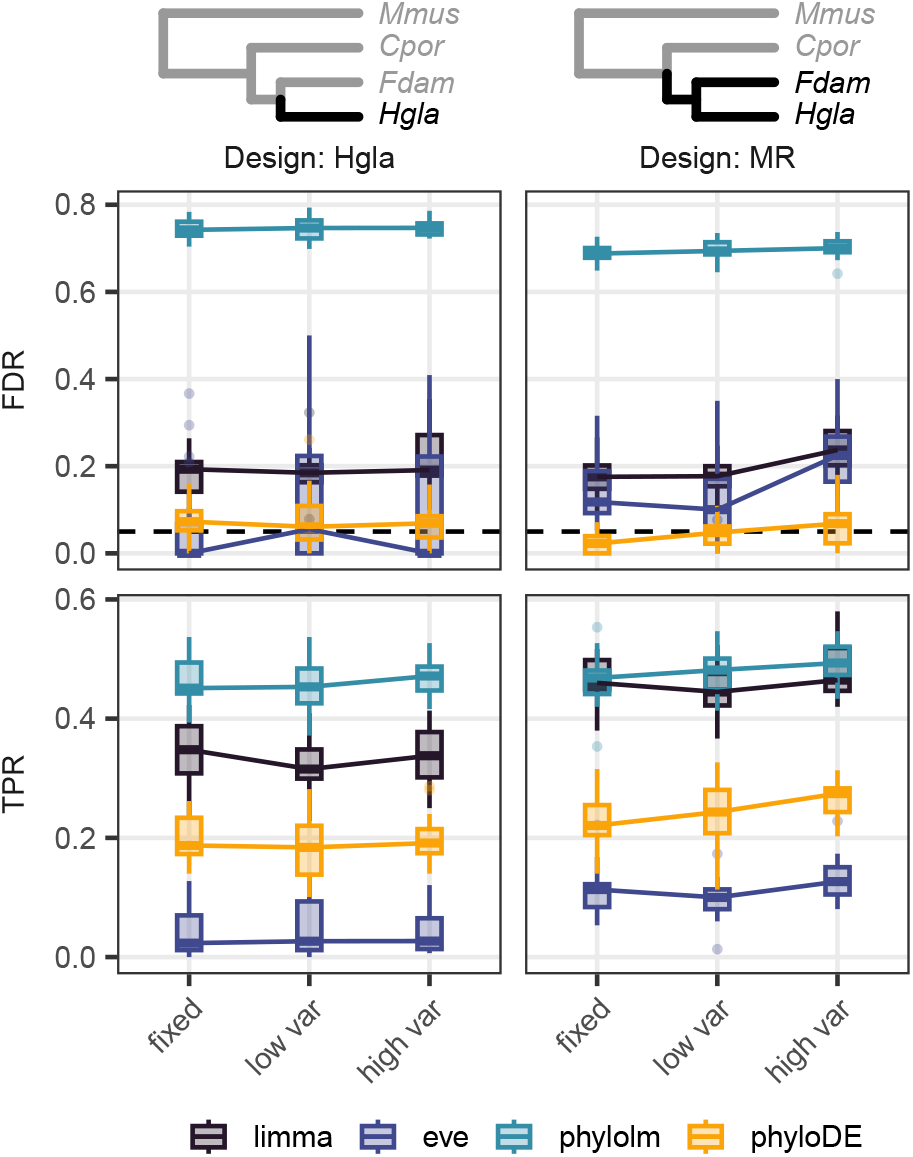
False Discovery Rate (FDR, top row) and True Positive Rate (TPR, bottom row) using Benjamini-Hochberg corrected p-values with a threshold of 5% (dashed horizontal line) for differential expression analysis between species *Hgla* versus the rest of the tree (Design: Hgla, left column), or both mole rat species versus the rest of the tree (Design: MR, right column), for 20 datasets simulated with the OU process using compcodeR on the empirical tree with 3 replicates per species, and using either empirical parameters for all genes (“fixed”), or varying parameters with a low or high variance (“low var” and “high var”), with a base effect size of 4. Methods are limma (with block correlation and empirical Bayes correction of fitted variances with trend), eve-ttt (with empirical p-values computed using 20 000 simulations under the null), phylolm (assuming a OU process with a fixed root), and phyloDE (assuming a OU process with a fixed root and empirical Bayes correction of fitted variances with trend). Our method phyloDE controls the FDR even under the high variance simulation setting.

**Figure S4:**
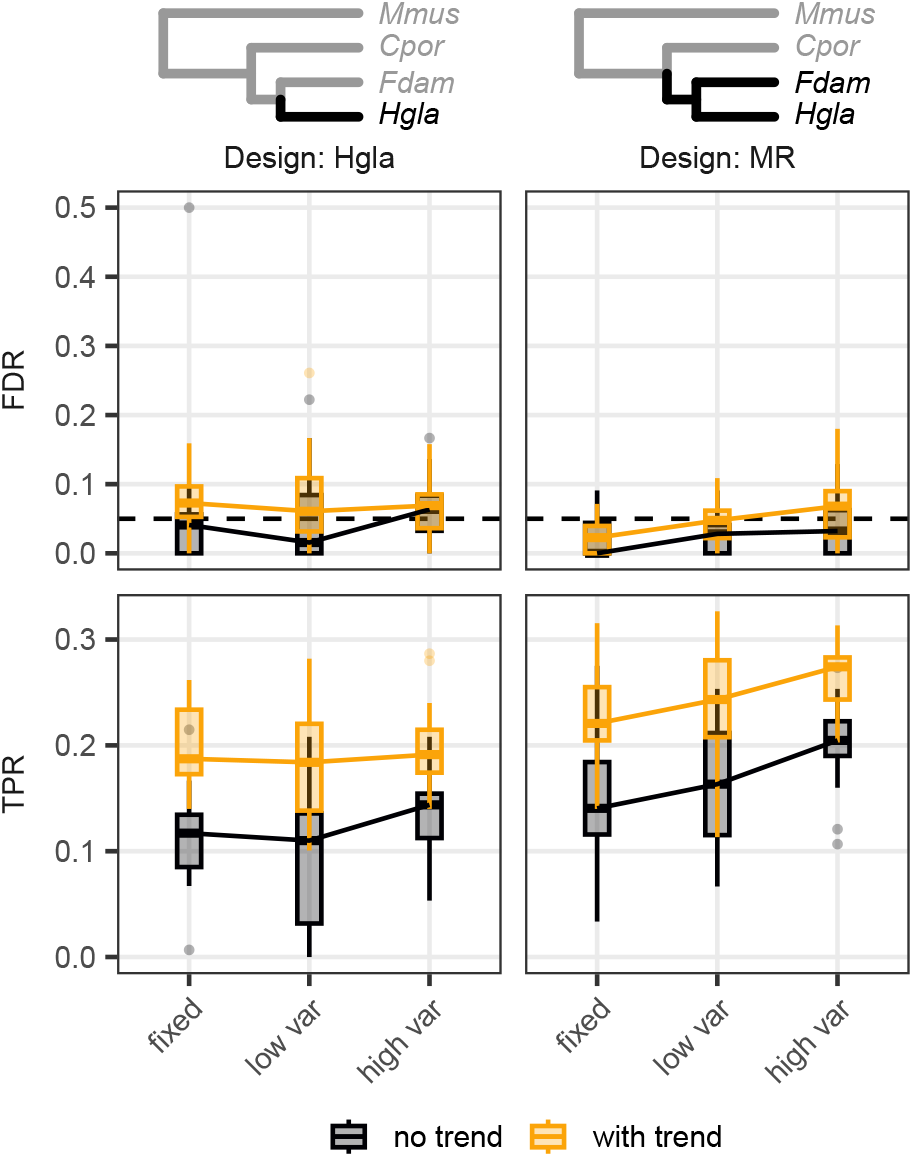
False Discovery Rate (FDR, top row) and True Positive Rate (TPR, bottom row) using Benjamini-Hochberg corrected p-values with a threshold of 5% (dashed horizontal line) for differential expression analysis between species *Hgla* versus the rest of the tree (Design: Hgla, left column), or both mole rat species versus the rest of the tree (Design: MR, right column), for 20 datasets simulated with the OU process using compcodeR on the empirical tree with 3 replicates per species, with fixed empirical parameters and a base effect size of 4. Comparison between phyloDE (assuming a OU process with a fixed root), with empirical Bayes correction, without or with the trend correction. Taking the mean-variance relationship slightly inflates in FDR, but substantially improves the TPR.

**Figure S5:**
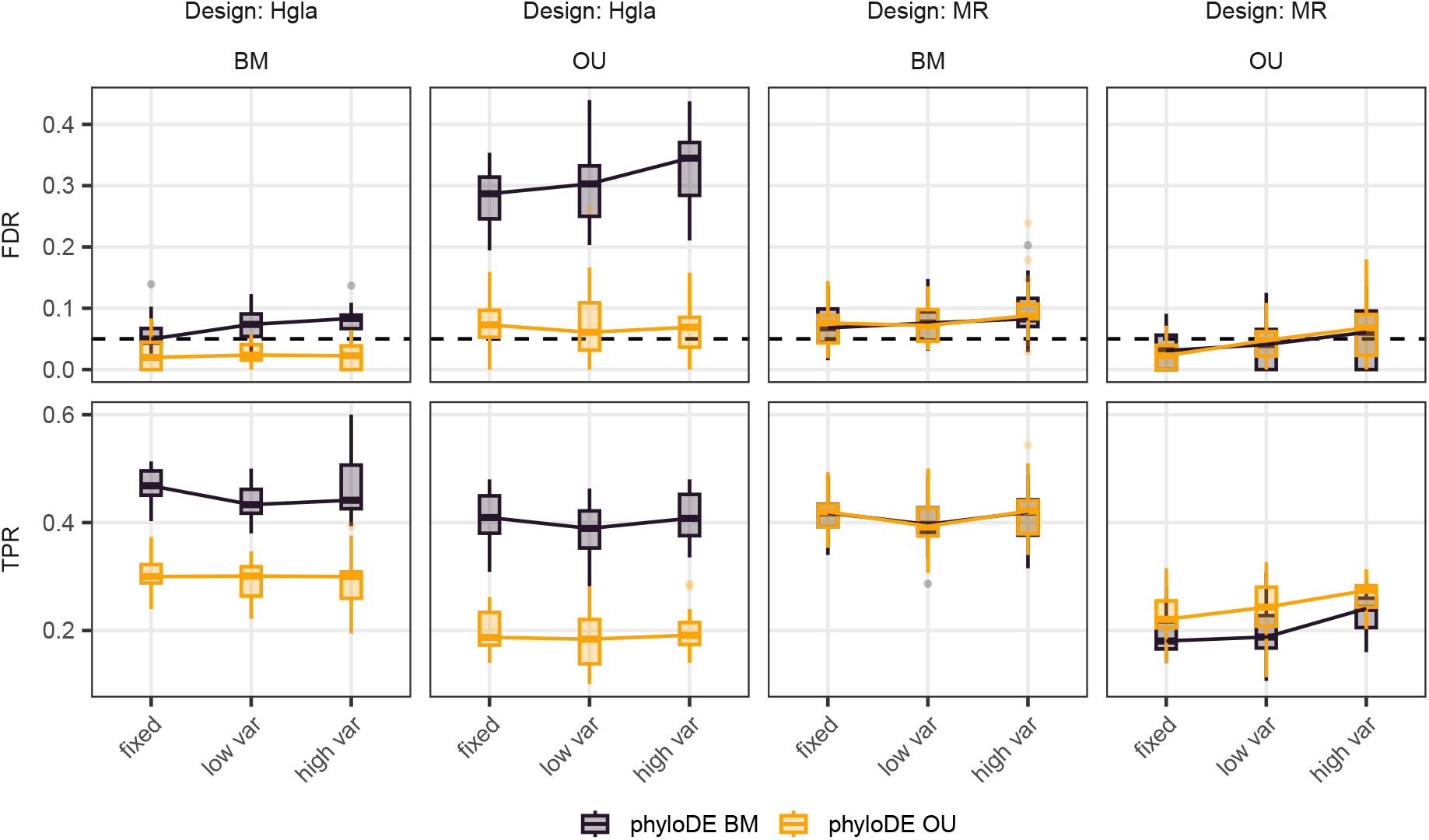
False Discovery Rate (FDR, top row) and True Positive Rate (TPR, bottom row) using Benjamini-Hochberg corrected p-values with a threshold of 5% (dashed horizontal line) for differential expression analysis between species *Hgla* versus the rest of the tree (Design: Hgla, left two columns), or both mole rat species versus the rest of the tree (Design: MR, right two columns), for 20 datasets simulated with the BM (left) or OU (right) processes using compcodeR on the empirical tree with 3 replicates per species, and using either empirical parameters for all genes (“fixed”), or varying parameters with a low or high variance (“low var” and “high var”). Methods are phyloDE, either assuming a BM process (dark grey) or a OU process (default, light orange), both with an empirical Bayes correction of fitted variances with trend. phyloDE assuming a OU process behaves better, even when the true simulating process is a BM.

## S3 Estimation of the parameters on a simple example

As described in the main text, we simulated a single dataset on the mole rat tree with three replicates per species, with 10 000 genes with no group effect, and each trait evolving according to a OU process with fixed selection strength *α* = 1.39, evolutionary rate variance 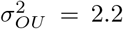, and withn-species variance *s*^2^ = 0.25, so that the total variance of Equation 4 was equal to *σ*^2^ = 1. We then applied evemodel, phylolm and phyloDE methods on this dataset, and compared the estimated parameters with the true values used for simulation.

**Figure S6:**
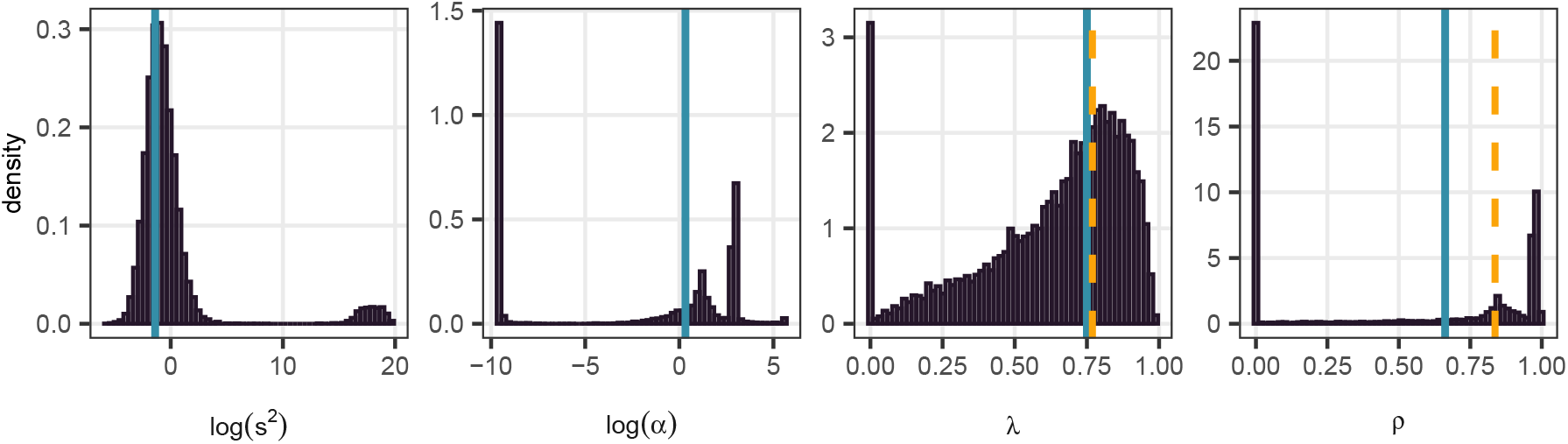
Estimation with phylolm and phyloDE of the parameters for one dataset with 10 000 independent genes following a OU process with fixed parameters on the mole rats tree with the three replicates per species. *s*^2^ and *α* are the original parameters used in phylolm (see Eq. 3), while *λ* and *ρ* are the transformed parameters that are regularized in phyloDE (see Eq. S5). The blue line shows the true parameter used in the simulation. The dashed light orange line shows the estimation using the trimmed mean 0.25, 0.05 on *λ* and *ρ* used by phyloDE.

**Figure S7:**
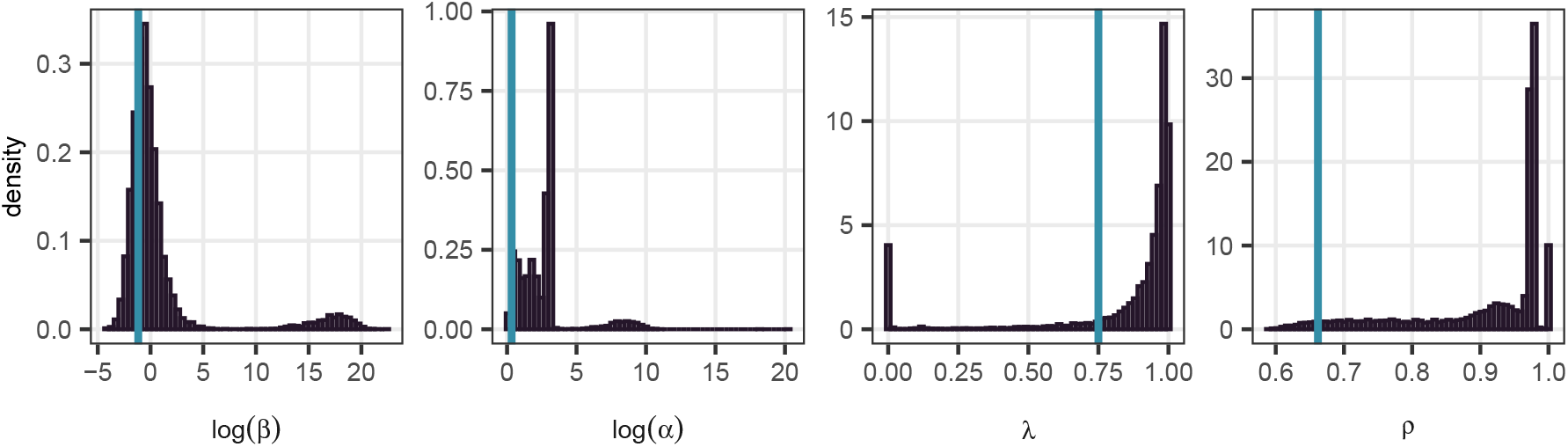
Histogram of estimated parameters using EVE (fonction fitOneTheta from package eve-model, Gillard *et al*., 2021), for 10 000 genes, all simulated with a OU process with fixed parameters on the mole rats tree with the three replicates per species. *β* and *α* are the original parameters used in EVE, while *λ* and *ρ* are the transformed parameters that are regularized in phyloDE, shown for comparison. The blue line shows the true parameter used in the simulation.

Fig. S6 shows the histograms of estimated variance parameters by phylolm and phyloDE. As in Fig. 1, these raw estimates are highly variable and un-reliable. Comparing Fig. S6 and S7, we can see that the transformed parameters *λ* and *ρ* estimated by phylolm and evemodel, that should be equal in theory, are quite different in practice. However, the maximized likelihood found by the two methods was very similar: over the 10 000 genes, the mean relative difference was equal to 0.01. This is expected, as the two methods do fit equivalent OU models on the data, although with different parametrizations, and the observed difference is likely due to numerical approximation errors, or different default specifications on the bounds of the parameters: for instance, evemodel allows extremelly hight values of *α*_*g*_ up to 6.1 *·* 10^8^, while phyloDE bounds this value at 265.7, which corresponds to a phylogenetic half life equal to 0.01% of the tree height.

This illustrates the fact that the likelihood surface can be very flat, and that the variance parameters are not correctly estimated on such small phylogenetic trees. While evemodel does not depend on the specific values of the parameters, as it uses a likelihood ratio test, the t-tests in phylolm do depend on these values, that will impact the effective sample size of the test through the correlation matrix. This makes the strong regularization of these parameters in phyloDE essencial to reduce the noise associated with the estimation of the variance parameters. In this exemple, we see that the trimmed mean produces reasonable estimates (Fig. S6, dashed lines), as it exploits the common structure across genes.

**Figure S8:**
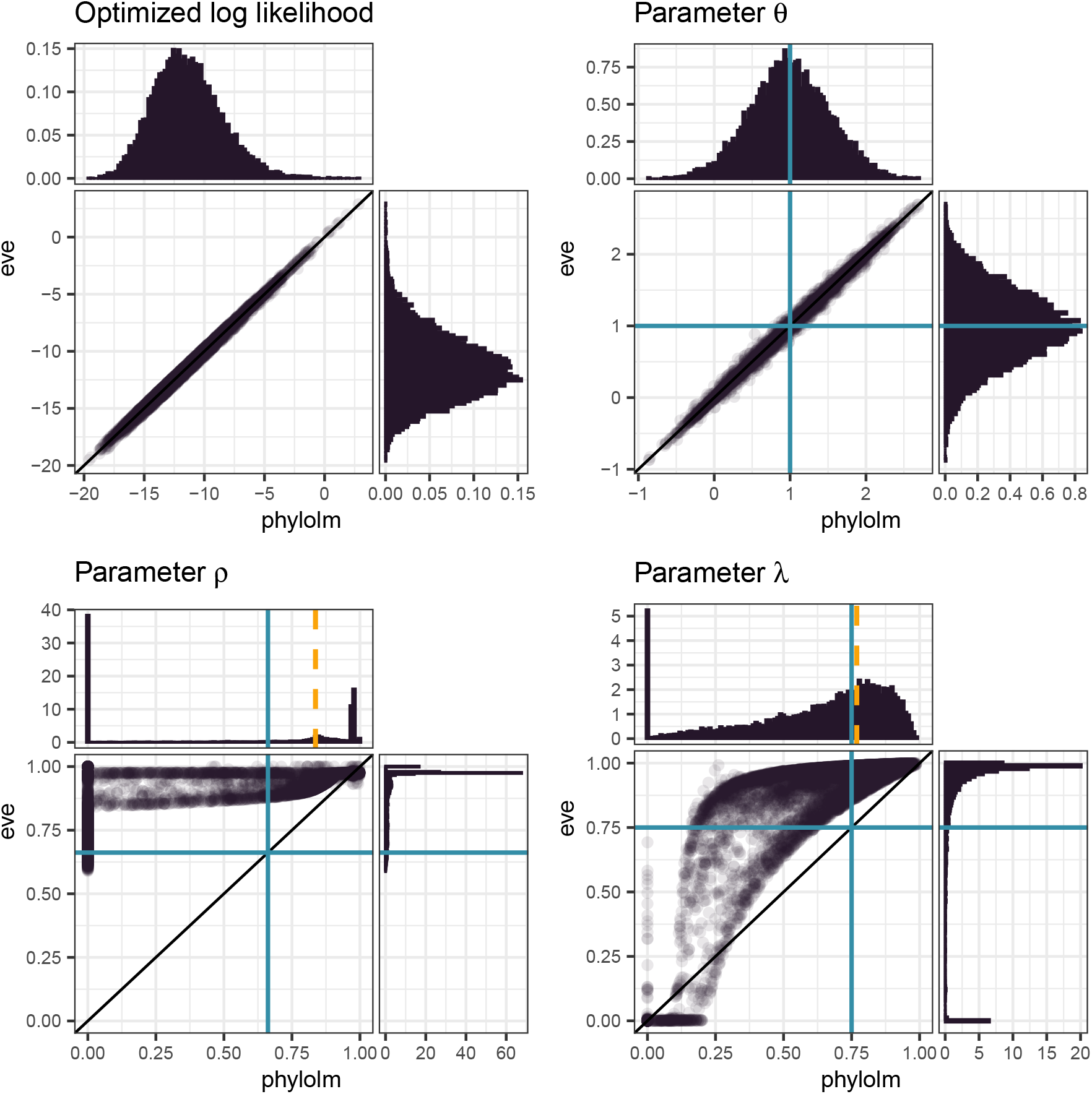
Comparison of the estimations with evemodel and phylolm of the normalized parameters for one dataset with 10 000 independent genes following a OU process with fixed parameters on the mole rats tree with the three replicates per species. *θ* is the common expectation to all species, and *λ* and *ρ* are the transformed parameters that are regularized in phyloDE (see Eq. S5). The blue line shows the true parameter used in the simulation. The dashed light orange line shows the estimation using the trimmed mean 0.25, 0.05 on *λ* and *ρ* used by phyloDE.

## S4 Behaviour of phyloDE with varying trim thresholds

**Figure S9:**
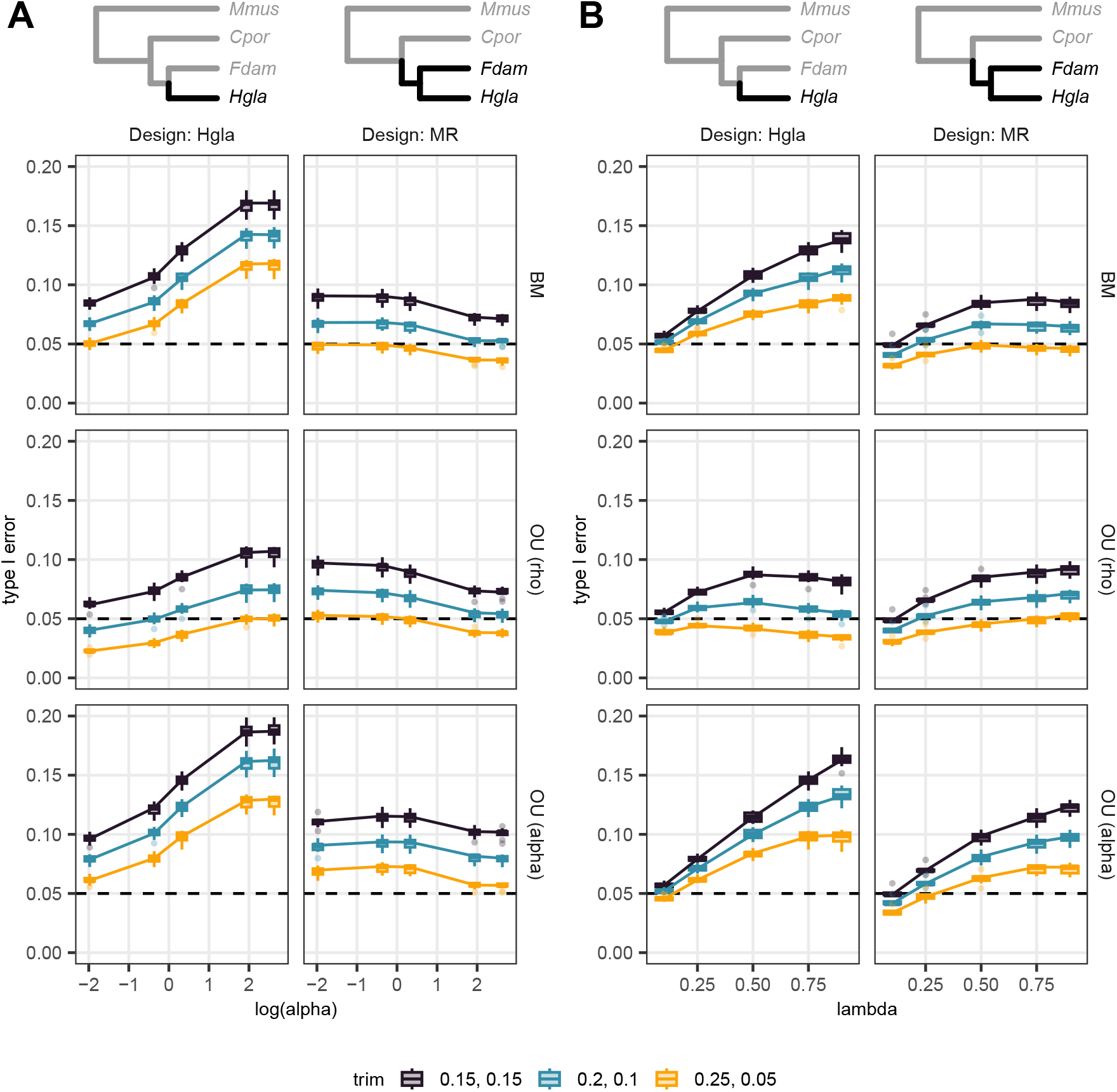
Type I error rate with a threshold of 5% (dashed horizontal line) for differential expression analysis between species *Hgla* versus the rest of the tree (Design: Hgla, left), or both mole rat species versus the rest of the tree (Design: MR, right), for 10 datasets simulated with the OU process with unit variance under the null hypothesis (no effect) for 10 000 genes on the empirical tree with 3 replicates per species. **A**: *λ* is fixed to 0.75, while *α* varies. **B**: log(2)*/α* is fixed to 0.5, while *λ* varies. We test three versions of phyloDE: assuming a BM and taking the trimmed mean of *λ* (first line); assuming a OU and taking the trimmed mean of *λ* and *ρ* (second line); and assuming a OU and taking the trimmed mean of *λ* and *α* (third line). Trim thresholds apply to transformed values. 30% of the values are trimmed in all cases. The *λ, ρ* parametrisation of the OU with trim values 0.25, 0.05 is the methods that has a type I error closest to the nominal value in all cases.

## S5 Lists of DE genes from the reanalysis of Daunesse *et al*. (2026)

We reporduce the list of genes for the two tissues (heart and liver) and the three designs (MR, hgla and fdam) found by phyloDE, using a threshold of 5% on the adjusted p-values. Genes are sorted by adjusted p-values and absolute value of the log-fold change.

**Table S1:** 96 DE genes found by phyloDE for the Heart data with the “MR” design.

| gene id | gene name | base log mean | log fold change | adjusted p-value (%) |
|---|---|---|---|---|
| ENSMUSG00000073639 | Rab18 | 7.31 | -7.15 | 0.00 |
| ENSMUSG00000033615 | Cplx1 | 0.45 | 6.42 | 0.07 |
| ENSMUSG00000030996 | Art1 | 7.55 | -5.70 | 0.07 |
| ENSMUSG00000027457 | Snph | 2.18 | 5.13 | 0.07 |
| ENSMUSG00000021668 | Polk | 4.04 | 2.14 | 0.07 |
| ENSMUSG00000056692 | Ilrun | 7.93 | -2.09 | 0.07 |
| ENSMUSG00000047658 | Gal3st3 | 5.74 | -5.51 | 0.12 |
| ENSMUSG00000033186 | Mzt1 | 6.18 | -5.36 | 0.12 |
| ENSMUSG00000036805 | Noxa1 | 0.34 | 4.95 | 0.12 |
| ENSMUSG00000022517 | Mgrn1 | 8.08 | -1.55 | 0.12 |
| ENSMUSG00000036322 | H2-Ea | 0.08 | 5.42 | 0.19 |
| ENSMUSG00000039662 | Icmt | 7.24 | -2.98 | 0.25 |
| ENSMUSG00000037894 | H2az1 | 4.99 | -4.62 | 0.35 |
| ENSMUSG00000020326 | Ccng1 | 9.06 | -2.75 | 0.35 |
| ENSMUSG00000035601 | Trmt10b | 3.80 | 1.84 | 0.50 |
| ENSMUSG00000024654 | Asrgl1 | 6.62 | -2.55 | 0.59 |
| ENSMUSG00000022594 | Lynx1 | 8.81 | -3.37 | 0.64 |
| ENSMUSG00000027165 | B230118H07Rik | 6.46 | -3.11 | 0.64 |
| ENSMUSG00000056666 | Retsat | 6.00 | 2.22 | 0.82 |
| ENSMUSG00000041020 | Map7d2 | 2.08 | 4.21 | 0.83 |
| ENSMUSG00000041248 | Kcnj1 | 0.13 | 4.11 | 0.89 |
| ENSMUSG00000024905 | Tesmin | 3.16 | -3.05 | 1.00 |
| ENSMUSG00000026058 | Khdrbs2 | 0.44 | 4.18 | 1.02 |
| ENSMUSG00000008999 | Bmp7 | 2.42 | 3.96 | 1.02 |
| ENSMUSG00000020062 | Slc5a8 | 0.50 | 3.73 | 1.02 |
| ENSMUSG00000011034 | Slc5a1 | 1.10 | 3.65 | 1.02 |
| ENSMUSG00000040726 | Hesx1 | 0.17 | 3.45 | 1.02 |
| ENSMUSG00000041079 | Rwdd2b | 4.47 | 1.68 | 1.02 |
| ENSMUSG00000006219 | Fblim1 | 7.67 | -2.83 | 1.25 |
| ENSMUSG00000027525 | Phactr3 | 1.24 | 4.49 | 1.31 |
| ENSMUSG00000024871 | Doc2g | 7.21 | -3.89 | 1.34 |
| ENSMUSG00000020630 | Rnaseh1 | 4.59 | 2.01 | 1.34 |
| ENSMUSG00000074227 | Spint2 | 5.05 | 2.78 | 1.50 |
| ENSMUSG00000020176 | Grb10 | 7.36 | -1.82 | 1.50 |
| ENSMUSG00000045636 | Mtus1 | 8.45 | -1.33 | 1.50 |
| ENSMUSG00000045291 | E330021D16Rik | 3.06 | 4.20 | 1.66 |
| ENSMUSG00000038173 | Enpp6 | 4.35 | -4.02 | 1.66 |
| ENSMUSG00000032216 | Nedd4 | 7.31 | -2.79 | 1.66 |
| ENSMUSG00000005968 | Tuft1 | 3.12 | 2.63 | 1.66 |
| ENSMUSG00000069255 | Dusp22 | 4.93 | 2.33 | 1.66 |
| ENSMUSG00000019843 | Fyn | 6.90 | -1.47 | 1.77 |
| ENSMUSG00000036912 | Piwil4 | 0.60 | 4.68 | 1.77 |
| ENSMUSG00000019785 | Clvs2 | 1.21 | 3.95 | 1.77 |
| ENSMUSG00000044086 | Lmod3 | 7.03 | -3.48 | 1.77 |
| ENSMUSG00000066975 | Cryba4 | 3.03 | -3.03 | 1.77 |
| ENSMUSG00000031026 | Trim66 | 1.17 | 2.91 | 1.77 |
| ENSMUSG00000022538 | Lsg1 | 4.71 | 1.56 | 1.77 |
| ENSMUSG00000024942 | Capn1 | 6.48 | 1.26 | 1.77 |
| ENSMUSG00000021890 | Eaf1 | 6.57 | -1.81 | 1.85 |
| ENSMUSG00000038205 | Prkab2 | 8.16 | -1.89 | 1.97 |
| ENSMUSG00000001249 | Hpn | 3.62 | 2.77 | 2.01 |
| ENSMUSG00000060450 | Rnf14 | 7.61 | -1.18 | 2.15 |
| ENSMUSG00000024780 | Cdc37l1 | 7.75 | -4.51 | 2.17 |
| ENSMUSG00000078938 | Syce3 | 0.27 | 4.49 | 2.21 |
| ENSMUSG00000015488 | Cacfd1 | 6.43 | -4.38 | 2.21 |
| ENSMUSG00000096210 | H1f0 | 6.81 | -3.93 | 2.22 |
| ENSMUSG00000031897 | Psmb10 | 6.28 | -2.67 | 2.22 |
| ENSMUSG00000043153 | Crppa | 4.07 | 1.74 | 2.30 |
| ENSMUSG00000025277 | Abhd6 | 4.23 | 2.62 | 2.31 |
| ENSMUSG00000009894 | Snap47 | 6.35 | -2.03 | 2.31 |
| ENSMUSG00000015568 | Lpl | 10.82 | -2.11 | 2.49 |
| ENSMUSG00000035805 | Mlc1 | 0.36 | 5.04 | 2.74 |
| ENSMUSG00000031637 | Lrp2bp | 4.85 | -4.47 | 2.78 |
| ENSMUSG00000031372 | Trex2 | 1.59 | 3.27 | 2.78 |
| ENSMUSG00000093661 | Eif4e3 | 5.83 | -2.07 | 2.95 |
| ENSMUSG00000007035 | Msh5 | 1.62 | 3.50 | 3.02 |
| ENSMUSG00000031709 | Tbc1d9 | 5.15 | 1.92 | 3.02 |
| ENSMUSG00000022390 | Zc3h7b | 7.76 | -1.33 | 3.02 |
| ENSMUSG00000022012 | Enox1 | 3.65 | 2.52 | 3.09 |
| ENSMUSG00000050272 | Dscam | 0.25 | 3.41 | 3.31 |
| ENSMUSG00000003161 | Sri | 6.75 | -1.45 | 3.31 |
| ENSMUSG00000031438 | Rnf128 | 6.27 | -2.57 | 3.45 |
| ENSMUSG00000032434 | Cmtm6 | 6.28 | -2.43 | 3.51 |
| ENSMUSG00000058743 | Kcnj14 | 4.33 | -3.84 | 3.60 |
| ENSMUSG00000042743 | Sgtb | 3.01 | 2.30 | 3.60 |
| ENSMUSG00000059995 | Atxn7l3 | 6.28 | -1.29 | 3.74 |
| ENSMUSG00000026778 | Prkcq | 5.22 | -4.21 | 3.74 |
| ENSMUSG00000050447 | Lypd6 | 1.03 | 3.00 | 3.74 |
| ENSMUSG00000032741 | Tpcn1 | 7.36 | 1.20 | 3.74 |
| ENSMUSG00000060002 | Chpt1 | 7.33 | -2.38 | 3.77 |
| ENSMUSG00000037348 | Paqr7 | 5.90 | -4.19 | 4.11 |
| ENSMUSG00000030168 | Adipor2 | 7.57 | 1.31 | 4.11 |
| ENSMUSG00000018931 | Natd1 | 6.22 | -2.29 | 4.39 |
| ENSMUSG00000002043 | Trappc6a | 4.45 | -3.45 | 4.58 |
| ENSMUSG00000024172 | St6gal2 | 0.60 | 3.04 | 4.58 |
| ENSMUSG00000052040 | Klf13 | 7.11 | -2.82 | 4.58 |
| ENSMUSG00000028271 | Gtf2b | 5.30 | 1.21 | 4.58 |
| ENSMUSG00000025867 | Cplx2 | 5.39 | -4.52 | 4.77 |
| ENSMUSG00000031775 | Pllp | 2.46 | 2.65 | 4.77 |
| ENSMUSG00000078607 | 1810010H24Rik | 1.39 | 2.86 | 4.91 |
| ENSMUSG00000047867 | Gimap6 | 5.95 | -3.29 | 4.96 |
| ENSMUSG00000031079 | Zfp300 | 2.68 | 2.60 | 4.96 |
| ENSMUSG00000024245 | Tmem178 | 2.43 | 2.57 | 4.96 |
| ENSMUSG00000020956 | Dtd2 | 4.83 | -2.35 | 4.96 |
| ENSMUSG00000019775 | Rgs17 | 2.11 | -2.08 | 4.96 |
| ENSMUSG00000024247 | Pkdcc | 6.03 | 1.23 | 4.96 |

**Table S2:** 157 DE genes found by phyloDE for the Heart data with the “hgla” design.

| gene id | gene name | base log mean | log fold change | adjusted p-value (%) |
|---|---|---|---|---|
| ENSMUSG00000022361 | Zhx1 | 6.07 | -6.07 | 0.00 |
| ENSMUSG00000042043 | Tbca | 6.14 | -6.14 | 0.00 |
| ENSMUSG00000029632 | Ndufa4 | 9.25 | -9.25 | 0.00 |
| ENSMUSG00000030265 | Kras | 6.47 | -6.47 | 0.00 |
| ENSMUSG00000096210 | H1f0 | 6.46 | -6.45 | 0.00 |
| ENSMUSG00000003970 | Rpl8 | 8.24 | -7.95 | 0.00 |
| ENSMUSG00000024780 | Cdc37l1 | 7.33 | -7.33 | 0.00 |
| ENSMUSG00000024287 | Thoc1 | 5.56 | -5.56 | 0.00 |
| ENSMUSG00000048100 | Taf13 | 5.50 | -5.50 | 0.00 |
| ENSMUSG00000049672 | Zbtb14 | 4.98 | -4.98 | 0.00 |
| ENSMUSG00000038067 | Csf3 | 0.00 | 4.57 | 0.00 |
| ENSMUSG00000041915 | Ammecr1l | 6.29 | -6.09 | 0.00 |
| ENSMUSG00000015749 | Anp32e | 6.64 | -6.64 | 0.00 |
| ENSMUSG00000032399 | Rpl4 | 8.86 | -8.53 | 0.00 |
| ENSMUSG00000027569 | Mrgbp | 4.71 | -4.71 | 0.00 |
| ENSMUSG00000032050 | Rdx | 7.28 | -7.07 | 0.00 |
| ENSMUSG00000045569 | Mc2r | 0.03 | 5.48 | 0.00 |
| ENSMUSG00000025816 | Sec61a2 | 5.69 | -5.69 | 0.00 |
| ENSMUSG00000014932 | Yes1 | 5.82 | -5.82 | 0.00 |
| ENSMUSG00000021270 | Hsp90aa1 | 8.66 | -8.16 | 0.00 |
| ENSMUSG00000022435 | Upk3a | 0.22 | 5.44 | 0.00 |
| ENSMUSG00000027067 | Ssrp1 | 6.66 | -6.44 | 0.00 |
| ENSMUSG00000003438 | Timm50 | 6.71 | -6.43 | 0.00 |
| ENSMUSG00000027556 | Car1 | 0.12 | 5.69 | 0.00 |
| ENSMUSG00000013663 | Pten | 6.81 | -6.73 | 0.00 |
| ENSMUSG00000020827 | Mink1 | 7.12 | -6.75 | 0.00 |
| ENSMUSG00000041959 | S100a10 | 6.15 | -5.24 | 0.00 |
| ENSMUSG00000055193 | Klk15 | 0.06 | 5.22 | 0.00 |
| ENSMUSG00000028266 | Lmo4 | 6.70 | -6.33 | 0.00 |
| ENSMUSG00000090862 | Rps13 | 7.37 | -7.37 | 0.00 |
| ENSMUSG00000037451 | Slc22a20 | 0.05 | 3.75 | 0.00 |
| ENSMUSG00000056153 | Socs6 | 5.97 | -5.83 | 0.00 |
| ENSMUSG00000026434 | Nucks1 | 6.85 | -6.85 | 0.00 |
| ENSMUSG00000032199 | Polr2m | 7.68 | -5.10 | 0.00 |
| ENSMUSG00000027956 | Tmem144 | 5.37 | -5.14 | 0.01 |
| ENSMUSG00000032249 | Anp32a | 7.28 | -6.33 | 0.01 |
| ENSMUSG00000002250 | Ppard | 6.33 | -2.58 | 0.01 |
| ENSMUSG00000037257 | Aagab | 5.57 | 2.28 | 0.01 |
| ENSMUSG00000015290 | Ubl4a | 7.30 | -2.95 | 0.02 |
| ENSMUSG00000036977 | Anapc10 | 4.35 | -4.20 | 0.02 |
| ENSMUSG00000049928 | Glp2r | 0.71 | 7.27 | 0.02 |
| ENSMUSG00000029203 | Ube2k | 6.62 | -4.95 | 0.03 |
| ENSMUSG00000036099 | Vezt | 6.12 | -3.59 | 0.03 |
| ENSMUSG00000048007 | Timm8a1 | 5.05 | -5.05 | 0.04 |
| ENSMUSG00000035765 | Dym | 6.58 | -3.83 | 0.05 |
| ENSMUSG00000047030 | Spata2 | 5.68 | -3.22 | 0.05 |
| ENSMUSG00000020333 | Acsl6 | 6.45 | -5.69 | 0.05 |
| ENSMUSG00000030350 | Prmt8 | 0.67 | 5.05 | 0.05 |
| ENSMUSG00000089809 | Rasgef1b | 4.41 | -4.14 | 0.06 |
| ENSMUSG00000023068 | Nus1 | 6.59 | -2.13 | 0.07 |
| ENSMUSG00000051978 | Erich1 | 4.12 | 2.21 | 0.07 |
| ENSMUSG00000036550 | Cnot1 | 7.79 | -3.92 | 0.08 |
| ENSMUSG00000047220 | Ccdc36 | 0.59 | 4.51 | 0.09 |
| ENSMUSG00000036333 | Kidins220 | 7.23 | 3.02 | 0.09 |
| ENSMUSG00000072964 | Bhlhb9 | 5.58 | -4.96 | 0.09 |
| ENSMUSG00000021096 | Ppm1a | 7.44 | -3.50 | 0.13 |
| ENSMUSG00000034189 | Hsdl1 | 6.17 | -1.93 | 0.15 |
| ENSMUSG00000038708 | Golga4 | 8.33 | -7.03 | 0.16 |
| ENSMUSG00000043441 | Gpr149 | 0.21 | 5.99 | 0.17 |
| ENSMUSG00000036805 | Noxa1 | 1.48 | 5.32 | 0.20 |
| ENSMUSG00000078653 | Cntd1 | 1.61 | 5.68 | 0.25 |
| ENSMUSG00000025265 | Fgd1 | 5.16 | -3.80 | 0.26 |
| ENSMUSG00000046345 | Smco1 | 4.61 | -3.62 | 0.27 |
| ENSMUSG00000094870 | Zfp131 | 5.50 | -1.86 | 0.27 |
| ENSMUSG00000025656 | Arhgef9 | 6.94 | -2.50 | 0.29 |
| ENSMUSG00000038736 | Nudcd1 | 5.14 | 2.80 | 0.32 |
| ENSMUSG00000018736 | Ndel1 | 6.07 | -5.53 | 0.36 |
| ENSMUSG00000044322 | Dsc1 | 0.16 | 4.93 | 0.36 |
| ENSMUSG00000020230 | Prmt2 | 5.22 | -4.69 | 0.42 |
| ENSMUSG00000050541 | Adra1b | 5.93 | -4.42 | 0.42 |
| ENSMUSG00000007850 | Hnrnph1 | 7.34 | -3.78 | 0.42 |
| ENSMUSG00000002279 | Lmf1 | 5.62 | 1.60 | 0.42 |
| ENSMUSG00000015217 | Hmgb3 | 5.63 | -4.28 | 0.44 |
| ENSMUSG00000024286 | Ccny | 7.33 | -1.53 | 0.46 |
| ENSMUSG00000020864 | Ankrd40 | 7.75 | -1.85 | 0.47 |
| ENSMUSG00000022010 | Tsc22d1 | 7.90 | -2.25 | 0.48 |
| ENSMUSG00000027744 | Stoml3 | 0.14 | 3.64 | 0.49 |
| ENSMUSG00000040842 | Szrd1 | 7.50 | -1.41 | 0.51 |
| ENSMUSG00000010663 | Fads1 | 5.98 | 1.58 | 0.53 |
| ENSMUSG00000034617 | Mtrr | 5.28 | -2.95 | 0.53 |
| ENSMUSG00000039194 | Rlbp1 | 0.43 | 5.05 | 0.55 |
| ENSMUSG00000053799 | Exoc6 | 5.82 | -2.74 | 0.55 |
| ENSMUSG00000038957 | Edc3 | 5.39 | -2.44 | 0.55 |
| ENSMUSG00000075569 | Rsph10b | 1.23 | 5.02 | 0.56 |
| ENSMUSG00000020107 | Anapc16 | 6.72 | -1.75 | 0.61 |
| ENSMUSG00000005886 | Ncoa2 | 6.98 | -5.84 | 0.61 |
| ENSMUSG00000041124 | Msantd4 | 6.18 | -1.78 | 0.64 |
| ENSMUSG00000051375 | Pcdh1 | 6.41 | -3.28 | 0.70 |
| ENSMUSG00000053399 | Adamts18 | 0.43 | 3.90 | 0.73 |
| ENSMUSG00000038845 | Phb | 8.19 | -7.09 | 0.74 |
| ENSMUSG00000027091 | Zc3h15 | 6.44 | -1.75 | 0.74 |
| ENSMUSG00000040848 | Sft2d2 | 6.26 | -1.92 | 0.78 |
| ENSMUSG00000033615 | Cplx1 | 2.09 | 6.27 | 0.79 |
| ENSMUSG00000039105 | Atp6v1g1 | 6.89 | -2.61 | 0.83 |
| ENSMUSG00000047786 | Lix1 | 5.81 | -3.61 | 0.88 |
| ENSMUSG00000026818 | Cel | 0.11 | 3.56 | 0.90 |
| ENSMUSG00000027332 | Ivd | 8.60 | -1.87 | 0.90 |
| ENSMUSG00000037020 | Wdr62 | 3.54 | 4.63 | 1.12 |
| ENSMUSG00000019832 | Rab32 | 3.81 | 2.30 | 1.23 |
| ENSMUSG00000043924 | Ncmap | 0.95 | 5.99 | 1.27 |
| ENSMUSG00000054999 | Naaladl1 | 0.44 | 5.52 | 1.41 |
| ENSMUSG00000004798 | Ulk2 | 7.14 | -2.31 | 1.48 |
| ENSMUSG00000031949 | Adat1 | 3.92 | -3.61 | 1.60 |
| ENSMUSG00000031897 | Psmb10 | 5.70 | -3.02 | 1.66 |
| ENSMUSG00000024327 | Slc39a7 | 6.76 | -3.24 | 1.68 |
| ENSMUSG00000030869 | Ndufab1 | 7.84 | -6.79 | 1.71 |
| ENSMUSG00000022594 | Lynx1 | 7.99 | -3.45 | 1.71 |
| ENSMUSG00000027134 | Lpcat4 | 3.78 | 3.08 | 1.71 |
| ENSMUSG00000032860 | P2ry2 | 6.06 | -2.12 | 1.72 |
| ENSMUSG00000051391 | Ywhag | 8.43 | -1.82 | 1.72 |
| ENSMUSG00000035984 | Nme5 | 3.50 | -3.50 | 1.95 |
| ENSMUSG00000026816 | Gtf3c5 | 5.45 | 1.37 | 2.16 |
| ENSMUSG00000079477 | Rab7 | 8.04 | -1.33 | 2.16 |
| ENSMUSG00000072640 | Lyrm9 | 6.31 | -2.52 | 2.17 |
| ENSMUSG00000040820 | Hlcs | 5.94 | -2.41 | 2.20 |
| ENSMUSG00000030559 | Rab38 | 1.31 | 5.55 | 2.33 |
| ENSMUSG00000060475 | Wtap | 6.28 | -3.95 | 2.33 |
| ENSMUSG00000019826 | Zbtb24 | 4.56 | 1.63 | 2.43 |
| ENSMUSG00000045094 | Arhgef37 | 5.53 | -2.98 | 2.56 |
| ENSMUSG00000054537 | Tmprss11e | 1.41 | 4.71 | 2.59 |
| ENSMUSG00000022015 | Tnfsf11 | 0.03 | 3.01 | 2.65 |
| ENSMUSG00000043300 | B3galnt1 | 4.84 | -4.36 | 2.67 |
| ENSMUSG00000044442 | N6amt1 | 5.44 | -2.33 | 2.68 |
| ENSMUSG00000029440 | Psmd9 | 5.72 | -2.16 | 2.70 |
| ENSMUSG00000040195 | Nemp1 | 4.38 | 2.05 | 2.72 |
| ENSMUSG00000038212 | Mfsd14b | 6.06 | -1.29 | 2.88 |
| ENSMUSG00000034007 | Scaper | 6.08 | -3.58 | 2.96 |
| ENSMUSG00000025758 | Plk4 | 4.32 | -3.90 | 3.00 |
| ENSMUSG00000028629 | Exo5 | 4.61 | -2.98 | 3.00 |
| ENSMUSG00000001802 | Lrp3 | 6.16 | -2.45 | 3.00 |
| ENSMUSG00000032475 | Nck1 | 6.16 | -1.64 | 3.00 |
| ENSMUSG00000006763 | Saal1 | 4.35 | 1.60 | 3.00 |
| ENSMUSG00000026327 | Serpinb11 | 0.07 | 4.52 | 3.08 |
| ENSMUSG00000023277 | Twf2 | 7.89 | -6.74 | 3.17 |
| ENSMUSG00000028223 | Decr1 | 8.07 | -2.22 | 3.18 |
| ENSMUSG00000056234 | Ncoa4 | 7.61 | -1.74 | 3.24 |
| ENSMUSG00000026443 | Lrrn2 | 4.47 | -4.11 | 3.30 |
| ENSMUSG00000027695 | Pld1 | 5.97 | 2.23 | 3.30 |
| ENSMUSG00000006649 | Nphs1 | 1.84 | 4.48 | 3.69 |
| ENSMUSG00000031669 | Gins3 | 3.51 | 4.55 | 3.80 |
| ENSMUSG00000025898 | Cwf19l2 | 4.54 | -4.42 | 3.84 |
| ENSMUSG00000053929 | Cyhr1 | 7.08 | -1.83 | 3.84 |
| ENSMUSG00000031562 | Dctd | 3.62 | 3.38 | 3.93 |
| ENSMUSG00000020657 | Dnajc27 | 5.54 | 1.89 | 4.05 |
| ENSMUSG00000028691 | Prdx1 | 8.15 | -2.53 | 4.11 |
| ENSMUSG00000033044 | Dhrs7c | 7.55 | 2.31 | 4.27 |
| ENSMUSG00000024726 | Carnmt1 | 6.92 | -1.72 | 4.28 |
| ENSMUSG00000049489 | Ccnq | 5.07 | -2.52 | 4.33 |
| ENSMUSG00000021619 | Atg10 | 4.92 | -2.06 | 4.46 |
| ENSMUSG00000022377 | Asap1 | 5.72 | 1.60 | 4.49 |
| ENSMUSG00000022372 | Sla | 3.65 | -2.97 | 4.58 |
| ENSMUSG00000033735 | Spr | 6.65 | -2.34 | 4.65 |
| ENSMUSG00000004933 | Matk | 1.73 | 4.67 | 4.81 |
| ENSMUSG00000024654 | Asrgl1 | 5.94 | -2.40 | 4.81 |
| ENSMUSG00000001569 | Nom1 | 5.51 | 1.83 | 4.81 |
| ENSMUSG00000041594 | Tmtc4 | 4.47 | 1.66 | 4.81 |
| ENSMUSG00000033416 | Gucd1 | 6.69 | -1.64 | 4.81 |

**Table S3:** 53 DE genes found by phyloDE for the Heart data with the “fdam” design.

| gene id | gene name | base log mean | log fold change | adjusted p-value (%) |
|---|---|---|---|---|
| ENSMUSG00000023456 | Tpi1 | 9.03 | -9.03 | 0.00 |
| ENSMUSG00000063410 | Stk24 | 6.96 | -6.96 | 0.00 |
| ENSMUSG00000029385 | Ccng2 | 5.71 | -5.71 | 0.00 |
| ENSMUSG00000015942 | Gtf2ird2 | 5.57 | -5.43 | 0.00 |
| ENSMUSG00000009900 | Wnt3a | 0.00 | 3.37 | 0.00 |
| ENSMUSG00000022197 | Pdzd2 | 7.56 | -7.43 | 0.01 |
| ENSMUSG00000024236 | Svil | 8.89 | -3.53 | 0.13 |
| ENSMUSG00000036958 | Cst11 | 0.00 | 6.12 | 0.16 |
| ENSMUSG00000024346 | Pfdn1 | 6.31 | -6.31 | 0.17 |
| ENSMUSG00000010080 | Epn3 | 6.17 | -5.79 | 0.17 |
| ENSMUSG00000070577 | Gm572 | 4.14 | -4.14 | 0.19 |
| ENSMUSG00000074591 | Ankub1 | 0.30 | 5.08 | 0.24 |
| ENSMUSG00000029651 | Mtus2 | 7.73 | -2.47 | 0.24 |
| ENSMUSG00000067081 | Asb18 | 5.47 | -5.47 | 0.32 |
| ENSMUSG00000030980 | Knop1 | 5.51 | -5.23 | 0.83 |
| ENSMUSG00000025217 | Btrc | 6.05 | 1.68 | 0.83 |
| ENSMUSG00000027443 | Cst12 | 0.24 | 5.69 | 0.95 |
| ENSMUSG00000079215 | Zfp664 | 7.20 | -2.27 | 1.29 |
| ENSMUSG00000064351 | mt-Co1 | 14.15 | -8.45 | 1.34 |
| ENSMUSG00000041293 | Adgrf1 | 1.25 | 6.34 | 1.52 |
| ENSMUSG00000045232 | Rln3 | 0.40 | 4.32 | 1.82 |
| ENSMUSG00000021556 | Golm1 | 6.38 | -2.76 | 1.82 |
| ENSMUSG00000031133 | Arhgef6 | 6.23 | 2.08 | 1.82 |
| ENSMUSG00000027879 | Sec22b | 6.16 | 1.57 | 1.86 |
| ENSMUSG00000030996 | Art1 | 6.12 | -5.67 | 1.94 |
| ENSMUSG00000045027 | Prss22 | 0.14 | 3.73 | 1.94 |
| ENSMUSG00000025239 | Limd1 | 6.95 | -2.07 | 1.94 |
| ENSMUSG00000042548 | Asxl1 | 6.53 | -4.74 | 2.18 |
| ENSMUSG00000029482 | Aacs | 4.98 | 2.47 | 2.18 |
| ENSMUSG00000020085 | Aifm2 | 6.43 | 2.38 | 2.18 |
| ENSMUSG00000053395 | Cacng8 | 0.32 | 5.92 | 2.76 |
| ENSMUSG00000044086 | Lmod3 | 6.30 | -4.03 | 2.76 |
| ENSMUSG00000027940 | Tpm3 | 6.83 | 3.25 | 2.76 |
| ENSMUSG00000048796 | Cyb561d1 | 5.20 | -2.69 | 2.76 |
| ENSMUSG00000038740 | Mvb12b | 6.81 | -2.22 | 2.76 |
| ENSMUSG00000045691 | Thtpa | 5.22 | 2.07 | 2.76 |
| ENSMUSG00000052214 | Opa3 | 7.04 | -3.06 | 3.07 |
| ENSMUSG00000044162 | Tnip3 | 2.70 | 4.40 | 3.14 |
| ENSMUSG00000024155 | Meiob | 0.35 | 3.61 | 3.14 |
| ENSMUSG00000032305 | Fam219b | 6.41 | -2.41 | 3.14 |
| ENSMUSG00000027430 | Dtd1 | 5.49 | 2.10 | 3.14 |
| ENSMUSG00000050144 | Slc25a44 | 5.58 | 1.48 | 3.14 |
| ENSMUSG00000036241 | Ube2r2 | 7.41 | -0.96 | 3.14 |
| ENSMUSG00000020923 | Ubtf | 7.03 | -1.66 | 3.24 |
| ENSMUSG00000006423 | C330007P06Rik | 5.41 | -2.70 | 3.33 |
| ENSMUSG00000032590 | Apeh | 6.78 | 1.51 | 3.88 |
| ENSMUSG00000027445 | Cst9 | 0.46 | 5.76 | 3.91 |
| ENSMUSG00000028217 | Cdh17 | 1.19 | 6.83 | 4.07 |
| ENSMUSG00000056222 | Spock1 | 2.26 | 4.17 | 4.07 |
| ENSMUSG00000031492 | Chrnb3 | 0.10 | 3.19 | 4.07 |
| ENSMUSG00000057948 | Unc13d | 3.30 | 3.18 | 4.07 |
| ENSMUSG00000031347 | Cetn2 | 5.75 | 2.19 | 4.07 |
| ENSMUSG00000041390 | Mdfic | 5.87 | -1.76 | 4.07 |

**Table S4:** 129 DE genes found by phyloDE for the Liver data with the “MR” design.

| gene id | gene name | base log mean | log fold change | adjusted p-value (%) |
|---|---|---|---|---|
| ENSMUSG00000073639 | Rab18 | 7.27 | -7.20 | 0.00 |
| ENSMUSG00000052974 | Cyp2f2 | 9.06 | -6.85 | 0.04 |
| ENSMUSG00000022505 | Emp2 | 3.75 | 3.52 | 0.04 |
| ENSMUSG00000032807 | Alox12b | 0.00 | 3.18 | 0.04 |
| ENSMUSG00000019891 | Dcbld1 | 4.30 | 2.45 | 0.07 |
| ENSMUSG00000051323 | Pcdh19 | 0.60 | 6.23 | 0.08 |
| ENSMUSG00000030307 | Slc6a11 | 0.79 | 7.25 | 0.09 |
| ENSMUSG00000033186 | Mzt1 | 5.47 | -4.69 | 0.09 |
| ENSMUSG00000050272 | Dscam | 0.00 | 3.29 | 0.09 |
| ENSMUSG00000030972 | Acsm5 | 7.81 | 2.41 | 0.09 |
| ENSMUSG00000054999 | Naaladl1 | 0.17 | 4.51 | 0.11 |
| ENSMUSG00000005089 | Slc1a2 | 8.19 | -6.66 | 0.15 |
| ENSMUSG00000028186 | Uox | 9.10 | -5.89 | 0.15 |
| ENSMUSG00000023978 | Prph2 | 0.00 | 5.14 | 0.15 |
| ENSMUSG00000046096 | Mosmo | 6.00 | -4.67 | 0.15 |
| ENSMUSG00000078938 | Syce3 | 0.11 | 4.44 | 0.15 |
| ENSMUSG00000032324 | Tspan3 | 5.58 | 3.21 | 0.15 |
| ENSMUSG00000056692 | Ilrun | 8.14 | -2.21 | 0.15 |
| ENSMUSG00000042096 | Dao | 2.88 | 5.93 | 0.16 |
| ENSMUSG00000036322 | H2-Ea | 0.09 | 5.72 | 0.16 |
| ENSMUSG00000039662 | Icmt | 6.46 | -2.03 | 0.24 |
| ENSMUSG00000020020 | Usp44 | 0.70 | 5.18 | 0.25 |
| ENSMUSG00000023094 | Msrb2 | 6.46 | 2.68 | 0.30 |
| ENSMUSG00000078584 | AU022252 | 7.03 | -2.50 | 0.34 |
| ENSMUSG00000029314 | Gpat3 | 3.02 | 4.01 | 0.39 |
| ENSMUSG00000033174 | Mgll | 7.40 | -3.31 | 0.53 |
| ENSMUSG00000012187 | Mogat1 | 0.89 | 5.73 | 0.54 |
| ENSMUSG00000045291 | E330021D16Rik | 2.25 | 4.62 | 0.54 |
| ENSMUSG00000063239 | Grm4 | 0.05 | 3.32 | 0.54 |
| ENSMUSG00000023963 | Cyp39a1 | 5.01 | 2.87 | 0.54 |
| ENSMUSG00000033096 | Apmap | 6.52 | 2.43 | 0.54 |
| ENSMUSG00000062312 | Erbb2 | 3.76 | 4.32 | 0.65 |
| ENSMUSG00000029471 | Camkk2 | 6.63 | -3.00 | 0.75 |
| ENSMUSG00000025494 | Sigirr | 6.08 | -3.00 | 0.75 |
| ENSMUSG00000024712 | Rfk | 7.07 | -2.30 | 0.75 |
| ENSMUSG00000029171 | Pgm2 | 4.94 | 2.14 | 0.80 |
| ENSMUSG00000069833 | Ahnak | 5.97 | 3.35 | 0.80 |
| ENSMUSG00000037224 | Zfyve28 | 0.48 | 4.60 | 1.03 |
| ENSMUSG00000012777 | Acp4 | 0.16 | 3.21 | 1.03 |
| ENSMUSG00000028862 | Map3k6 | 3.41 | 2.50 | 1.03 |
| ENSMUSG00000015652 | Steap1 | 1.65 | 3.37 | 1.11 |
| ENSMUSG00000036943 | Rab8b | 4.87 | 2.28 | 1.11 |
| ENSMUSG00000024601 | Isoc1 | 7.22 | -1.80 | 1.20 |
| ENSMUSG00000099517 | H3c8 | 2.94 | 3.74 | 1.31 |
| ENSMUSG00000028185 | Dnase2b | 5.85 | -4.12 | 1.44 |
| ENSMUSG00000056162 | Cndp1 | 0.86 | 5.35 | 1.45 |
| ENSMUSG00000029123 | Stk32b | 0.05 | 4.95 | 1.45 |
| ENSMUSG00000064307 | Lrrc51 | 3.03 | 2.62 | 1.45 |
| ENSMUSG00000024130 | Abca3 | 7.83 | -1.50 | 1.45 |
| ENSMUSG00000049404 | Rarres1 | 7.25 | -4.61 | 1.49 |
| ENSMUSG00000020007 | Il20ra | 0.04 | 4.08 | 1.49 |
| ENSMUSG00000020288 | Ahsa2 | 6.08 | -3.22 | 1.49 |
| ENSMUSG00000037894 | H2az1 | 5.11 | -4.82 | 1.50 |
| ENSMUSG00000010066 | Cacna2d2 | 0.21 | 4.22 | 1.51 |
| ENSMUSG00000025726 | Slc28a1 | 0.60 | 4.07 | 1.54 |
| ENSMUSG00000028885 | Smpdl3b | 3.38 | 4.18 | 1.65 |
| ENSMUSG00000057722 | Lepr | 2.23 | 5.32 | 1.69 |
| ENSMUSG00000025758 | Plk4 | 4.75 | -3.57 | 1.69 |
| ENSMUSG00000021583 | Erap1 | 7.63 | -1.58 | 1.69 |
| ENSMUSG00000058589 | Anks1b | 0.99 | 4.54 | 1.73 |
| ENSMUSG00000030905 | Crym | 2.81 | 4.82 | 1.83 |
| ENSMUSG00000043487 | Acot6 | 2.27 | 4.04 | 1.83 |
| ENSMUSG00000001039 | B9d1 | 2.70 | 3.15 | 1.83 |
| ENSMUSG00000020329 | Polrmt | 6.21 | 1.67 | 1.83 |
| ENSMUSG00000046613 | Vwa5b2 | 0.08 | 3.64 | 1.90 |
| ENSMUSG00000027762 | Sucnr1 | 5.97 | -4.66 | 2.10 |
| ENSMUSG00000022895 | Ets2 | 5.95 | 1.99 | 2.17 |
| ENSMUSG00000032050 | Rdx | 8.07 | -4.66 | 2.39 |
| ENSMUSG00000020142 | Slc1a4 | 3.50 | 3.81 | 2.39 |
| ENSMUSG00000039058 | Ak5 | 0.39 | 3.80 | 2.39 |
| ENSMUSG00000043843 | Tmem145 | 0.16 | 3.42 | 2.39 |
| ENSMUSG00000061046 | Haghl | 4.00 | -3.14 | 2.39 |
| ENSMUSG00000020838 | Slc6a4 | 2.55 | 3.03 | 2.39 |
| ENSMUSG00000027035 | Cers6 | 4.39 | 2.69 | 2.39 |
| ENSMUSG00000025355 | Mmp19 | 7.29 | -2.55 | 2.39 |
| ENSMUSG00000019832 | Rab32 | 6.35 | 1.69 | 2.39 |
| ENSMUSG00000021996 | Esd | 8.10 | -1.45 | 2.39 |
| ENSMUSG00000015405 | Ace2 | 2.24 | 4.49 | 2.40 |
| ENSMUSG00000048232 | Fbxo10 | 3.03 | 2.03 | 2.40 |
| ENSMUSG00000074505 | Fat3 | 0.22 | 3.63 | 2.41 |
| ENSMUSG00000027489 | Necab3 | 0.58 | 4.13 | 2.51 |
| ENSMUSG00000041187 | Prkd2 | 5.17 | 2.36 | 2.52 |
| ENSMUSG00000047656 | Trpt1 | 3.53 | 2.84 | 2.56 |
| ENSMUSG00000039530 | Tusc3 | 5.53 | -3.71 | 2.73 |
| ENSMUSG00000079554 | Aox2 | 0.69 | 3.42 | 2.73 |
| ENSMUSG00000013338 | Fer1l4 | 0.00 | 3.33 | 2.82 |
| ENSMUSG00000019718 | L3hypdh | 6.18 | -1.95 | 2.87 |
| ENSMUSG00000033053 | 1700028P14Rik | 0.00 | 2.54 | 2.89 |
| ENSMUSG00000029669 | Tspan12 | 8.04 | -2.39 | 2.89 |
| ENSMUSG00000028294 | Cfap206 | 0.18 | 3.54 | 3.03 |
| ENSMUSG00000055660 | Mettl4 | 5.65 | -1.79 | 3.19 |
| ENSMUSG00000010609 | Psen2 | 8.45 | -2.77 | 3.35 |
| ENSMUSG00000044442 | N6amt1 | 5.24 | -1.77 | 3.40 |
| ENSMUSG00000027015 | Cybrd1 | 3.34 | 3.28 | 3.54 |
| ENSMUSG00000029638 | Glcci1 | 5.58 | -3.16 | 3.54 |
| ENSMUSG00000004902 | Slc25a18 | 0.77 | 4.91 | 3.54 |
| ENSMUSG00000028649 | Macf1 | 8.76 | -3.57 | 3.54 |
| ENSMUSG00000069378 | Prdm6 | 0.97 | 4.12 | 3.59 |
| ENSMUSG00000045967 | Gpr158 | 2.44 | 3.99 | 3.59 |
| ENSMUSG00000051703 | Tmem198 | 0.32 | 3.85 | 3.59 |
| ENSMUSG00000044813 | Shb | 5.95 | -3.22 | 3.70 |
| ENSMUSG00000032434 | Cmtm6 | 7.88 | -2.25 | 3.70 |
| ENSMUSG00000032788 | Pdxk | 7.17 | -1.52 | 3.70 |
| ENSMUSG00000029032 | Arhgef16 | 3.50 | 2.70 | 3.71 |
| ENSMUSG00000026870 | Cutal | 5.71 | -4.08 | 3.83 |
| ENSMUSG00000029458 | Brap | 7.48 | -1.43 | 3.83 |
| ENSMUSG00000006056 | Calcoco2 | 2.81 | 3.98 | 3.87 |
| ENSMUSG00000017009 | Sdc4 | 8.56 | -2.51 | 3.87 |
| ENSMUSG00000020630 | Rnaseh1 | 4.69 | 1.49 | 3.87 |
| ENSMUSG00000032673 | Prorsd1 | 5.22 | -2.18 | 3.88 |
| ENSMUSG00000024287 | Thoc1 | 5.88 | -3.48 | 3.89 |
| ENSMUSG00000022179 | 4931414P19Rik | 4.13 | 1.56 | 3.89 |
| ENSMUSG00000067878 | Map7d3 | 2.53 | 3.95 | 4.13 |
| ENSMUSG00000022751 | Nit2 | 7.81 | -1.83 | 4.46 |
| ENSMUSG00000036805 | Noxa1 | 1.13 | 4.74 | 4.52 |
| ENSMUSG00000025736 | Jmjd8 | 6.13 | -1.72 | 4.52 |
| ENSMUSG00000003623 | Crot | 8.89 | -2.24 | 4.59 |
| ENSMUSG00000050822 | Slc29a4 | 0.31 | 3.42 | 4.59 |
| ENSMUSG00000027698 | Nceh1 | 6.42 | -2.93 | 4.67 |
| ENSMUSG00000030315 | Vgll4 | 6.01 | -2.18 | 4.67 |
| ENSMUSG00000058441 | Panx2 | 1.12 | 3.59 | 4.69 |
| ENSMUSG00000034391 | Fbxo15 | 1.57 | 3.29 | 4.69 |
| ENSMUSG00000022200 | Golph3 | 7.41 | -1.40 | 4.69 |
| ENSMUSG00000051950 | B3glct | 4.46 | 1.66 | 4.78 |
| ENSMUSG00000026421 | Csrp1 | 5.88 | 2.35 | 4.78 |
| ENSMUSG00000004530 | Coro1c | 5.87 | 1.13 | 4.83 |
| ENSMUSG00000003476 | Crhr2 | 0.50 | 3.79 | 4.88 |
| ENSMUSG00000032849 | Abcc4 | 4.45 | 2.84 | 4.88 |
| ENSMUSG00000058886 | Deaf1 | 4.93 | 1.91 | 4.90 |

**Table S5:** 131 DE genes found by phyloDE for the Liver data with the “hgla” design.

| gene id | gene name | base log mean | log fold change | adjusted p-value (%) |
|---|---|---|---|---|
| ENSMUSG00000003970 | Rpl8 | 8.18 | -8.18 | 0.00 |
| ENSMUSG00000055193 | Klk15 | 0.00 | 5.01 | 0.00 |
| ENSMUSG00000032050 | Rdx | 7.67 | -7.67 | 0.00 |
| ENSMUSG00000027518 | 1700021F07Rik | 0.00 | 5.39 | 0.00 |
| ENSMUSG00000014932 | Yes1 | 6.73 | -6.73 | 0.00 |
| ENSMUSG00000029632 | Ndufa4 | 7.85 | -7.85 | 0.00 |
| ENSMUSG00000026434 | Nucks1 | 7.14 | -7.07 | 0.00 |
| ENSMUSG00000035171 | 1110059E24Rik | 5.20 | -5.20 | 0.00 |
| ENSMUSG00000005886 | Ncoa2 | 7.59 | -7.52 | 0.00 |
| ENSMUSG00000032399 | Rpl4 | 8.66 | -8.38 | 0.00 |
| ENSMUSG00000030265 | Kras | 6.64 | -6.64 | 0.00 |
| ENSMUSG00000056153 | Socs6 | 6.24 | -6.24 | 0.00 |
| ENSMUSG00000049672 | Zbtb14 | 4.91 | -4.91 | 0.00 |
| ENSMUSG00000042043 | Tbca | 6.08 | -6.08 | 0.00 |
| ENSMUSG00000090862 | Rps13 | 6.88 | -6.88 | 0.00 |
| ENSMUSG00000013663 | Pten | 8.00 | -7.80 | 0.00 |
| ENSMUSG00000096210 | H1f0 | 6.64 | -6.64 | 0.00 |
| ENSMUSG00000022361 | Zhx1 | 6.76 | -6.64 | 0.00 |
| ENSMUSG00000021270 | Hsp90aa1 | 9.13 | -8.23 | 0.00 |
| ENSMUSG00000027067 | Ssrp1 | 6.73 | -6.52 | 0.00 |
| ENSMUSG00000024780 | Cdc37l1 | 6.43 | -6.33 | 0.00 |
| ENSMUSG00000037816 | Fbxw17 | 4.08 | -4.08 | 0.00 |
| ENSMUSG00000037159 | Wee2 | 0.00 | 3.89 | 0.00 |
| ENSMUSG00000024287 | Thoc1 | 5.50 | -5.43 | 0.00 |
| ENSMUSG00000027956 | Tmem144 | 6.10 | -6.10 | 0.00 |
| ENSMUSG00000026285 | Pdcd1 | 0.24 | 6.82 | 0.00 |
| ENSMUSG00000048100 | Taf13 | 5.54 | -5.54 | 0.00 |
| ENSMUSG00000036095 | Dgkb | 0.13 | 5.20 | 0.00 |
| ENSMUSG00000024327 | Slc39a7 | 7.55 | -4.21 | 0.00 |
| ENSMUSG00000020827 | Mink1 | 7.07 | -6.68 | 0.00 |
| ENSMUSG00000028266 | Lmo4 | 5.76 | -5.76 | 0.00 |
| ENSMUSG00000074003 | Gucy2d | 0.00 | 3.39 | 0.00 |
| ENSMUSG00000003438 | Timm50 | 6.15 | -5.82 | 0.00 |
| ENSMUSG00000030869 | Ndufab1 | 6.28 | -6.08 | 0.00 |
| ENSMUSG00000040472 | Rabggta | 5.86 | -3.39 | 0.01 |
| ENSMUSG00000032249 | Anp32a | 6.98 | -5.79 | 0.01 |
| ENSMUSG00000025089 | Gfra1 | 7.60 | -4.42 | 0.01 |
| ENSMUSG00000044172 | Ptx4 | 0.21 | 5.50 | 0.01 |
| ENSMUSG00000022454 | Nell2 | 0.18 | 3.84 | 0.01 |
| ENSMUSG00000030905 | Crym | 3.57 | 6.57 | 0.01 |
| ENSMUSG00000068327 | Tlx2 | 0.13 | 5.65 | 0.01 |
| ENSMUSG00000042256 | Ptchd4 | 0.08 | 5.27 | 0.01 |
| ENSMUSG00000041915 | Ammecr1l | 6.48 | -6.06 | 0.02 |
| ENSMUSG00000036977 | Anapc10 | 4.31 | -4.31 | 0.04 |
| ENSMUSG00000036550 | Cnot1 | 8.21 | -3.59 | 0.04 |
| ENSMUSG00000021098 | 4930447C04Rik | 0.00 | 3.37 | 0.04 |
| ENSMUSG00000018736 | Ndel1 | 5.60 | -5.24 | 0.04 |
| ENSMUSG00000038845 | Phb | 7.46 | -7.14 | 0.05 |
| ENSMUSG00000036099 | Vezt | 5.92 | -4.68 | 0.05 |
| ENSMUSG00000022010 | Tsc22d1 | 7.33 | -4.32 | 0.05 |
| ENSMUSG00000038990 | Cables2 | 4.92 | -4.65 | 0.06 |
| ENSMUSG00000027104 | Atf2 | 6.43 | -3.59 | 0.07 |
| ENSMUSG00000060475 | Wtap | 6.18 | -3.84 | 0.09 |
| ENSMUSG00000015709 | Arnt2 | 1.54 | 4.49 | 0.09 |
| ENSMUSG00000015749 | Anp32e | 5.71 | -5.64 | 0.11 |
| ENSMUSG00000031402 | Mpp1 | 5.89 | 1.95 | 0.16 |
| ENSMUSG00000029334 | Prkg2 | 0.68 | 5.87 | 0.17 |
| ENSMUSG00000056155 | Nanos3 | 0.08 | 4.28 | 0.17 |
| ENSMUSG00000059857 | Ntng1 | 0.03 | 3.21 | 0.19 |
| ENSMUSG00000070498 | Tmem132b | 0.17 | 5.28 | 0.19 |
| ENSMUSG00000027569 | Mrgbp | 4.14 | -4.06 | 0.19 |
| ENSMUSG00000024578 | Il17b | 0.88 | 6.41 | 0.21 |
| ENSMUSG00000051375 | Pcdh1 | 7.43 | -3.55 | 0.26 |
| ENSMUSG00000030352 | Tspan9 | 6.96 | -1.79 | 0.29 |
| ENSMUSG00000078653 | Cntd1 | 0.69 | 6.26 | 0.36 |
| ENSMUSG00000044499 | Hs3st5 | 0.20 | 5.44 | 0.37 |
| ENSMUSG00000031250 | Tnmd | 0.22 | 4.25 | 0.38 |
| ENSMUSG00000021096 | Ppm1a | 7.67 | -5.92 | 0.46 |
| ENSMUSG00000022483 | Col2a1 | 0.47 | 4.97 | 0.46 |
| ENSMUSG00000015702 | Anxa9 | 2.05 | 4.90 | 0.53 |
| ENSMUSG00000025898 | Cwf19l2 | 4.84 | -4.84 | 0.54 |
| ENSMUSG00000023122 | Sult1c2 | 1.33 | 5.11 | 0.54 |
| ENSMUSG00000051367 | Six1 | 0.11 | 3.59 | 0.54 |
| ENSMUSG00000052372 | Il1rapl1 | 0.14 | 4.07 | 0.58 |
| ENSMUSG00000022419 | Deptor | 7.58 | -3.41 | 0.62 |
| ENSMUSG00000046096 | Mosmo | 4.89 | -4.89 | 0.72 |
| ENSMUSG00000028030 | Tbck | 6.29 | -2.97 | 0.83 |
| ENSMUSG00000028294 | Cfap206 | 0.85 | 4.37 | 0.83 |
| ENSMUSG00000030963 | Umod | 0.24 | 6.12 | 0.83 |
| ENSMUSG00000025081 | Tdrd1 | 0.63 | 5.30 | 0.83 |
| ENSMUSG00000028427 | Aqp7 | 1.41 | 4.82 | 0.86 |
| ENSMUSG00000079554 | Aox2 | 1.35 | 4.15 | 0.98 |
| ENSMUSG00000026621 | Mtarc1 | 8.36 | -2.47 | 1.01 |
| ENSMUSG00000020173 | Cobl | 6.16 | -3.87 | 1.10 |
| ENSMUSG00000022613 | Miox | 1.89 | 7.48 | 1.11 |
| ENSMUSG00000038148 | Cldn16 | 0.05 | 2.73 | 1.25 |
| ENSMUSG00000037260 | Hgsnat | 6.81 | -1.60 | 1.29 |
| ENSMUSG00000045394 | Epcam | 3.76 | 4.05 | 1.48 |
| ENSMUSG00000020288 | Ahsa2 | 5.39 | -3.64 | 1.48 |
| ENSMUSG00000055333 | Fat2 | 1.04 | 5.67 | 1.55 |
| ENSMUSG00000048007 | Timm8a1 | 5.27 | -5.20 | 1.55 |
| ENSMUSG00000031844 | Hsd17b2 | 9.19 | -5.64 | 1.57 |
| ENSMUSG00000053166 | Cdh22 | 0.34 | 5.17 | 1.57 |
| ENSMUSG00000047030 | Spata2 | 4.87 | -3.64 | 1.57 |
| ENSMUSG00000035310 | Lin54 | 5.32 | -3.02 | 1.64 |
| ENSMUSG00000003354 | Ccdc65 | 0.16 | 2.83 | 1.64 |
| ENSMUSG00000019900 | Rfx6 | 0.00 | 2.65 | 1.64 |
| ENSMUSG00000034219 | Septin14 | 0.29 | 3.88 | 1.68 |
| ENSMUSG00000038544 | Inip | 4.93 | 1.70 | 1.71 |
| ENSMUSG00000026433 | Rab29 | 3.67 | 3.23 | 1.75 |
| ENSMUSG00000040037 | Negr1 | 0.89 | 4.87 | 1.84 |
| ENSMUSG00000025816 | Sec61a2 | 4.22 | -4.11 | 1.91 |
| ENSMUSG00000025485 | Ric8a | 6.13 | 1.51 | 1.91 |
| ENSMUSG00000059495 | Arhgef12 | 8.23 | -4.49 | 1.95 |
| ENSMUSG00000032454 | Rbp2 | 0.00 | 3.17 | 1.97 |
| ENSMUSG00000039715 | Wdr34 | 4.44 | 1.99 | 2.81 |
| ENSMUSG00000045573 | Penk | 0.20 | 4.06 | 2.97 |
| ENSMUSG00000021951 | Eef1akmt1 | 4.32 | 2.20 | 2.97 |
| ENSMUSG00000089809 | Rasgef1b | 5.80 | -5.28 | 3.18 |
| ENSMUSG00000024654 | Asrgl1 | 5.31 | -4.20 | 3.25 |
| ENSMUSG00000039395 | Mreg | 4.94 | -4.64 | 3.36 |
| ENSMUSG00000036943 | Rab8b | 5.40 | 2.40 | 3.36 |
| ENSMUSG00000028649 | Macf1 | 8.01 | -4.10 | 3.47 |
| ENSMUSG00000025758 | Plk4 | 3.94 | -3.87 | 3.47 |
| ENSMUSG00000037451 | Slc22a20 | 0.23 | 2.93 | 3.81 |
| ENSMUSG00000045282 | Tmem86b | 6.41 | -4.05 | 3.84 |
| ENSMUSG00000029622 | Arpc1b | 6.42 | 1.87 | 3.91 |
| ENSMUSG00000021000 | Mia2 | 8.54 | -1.69 | 3.91 |
| ENSMUSG00000038775 | Vill | 4.03 | -4.03 | 4.02 |
| ENSMUSG00000032324 | Tspan3 | 6.42 | 3.05 | 4.03 |
| ENSMUSG00000039717 | Ralyl | 0.00 | 2.31 | 4.09 |
| ENSMUSG00000006411 | Nectin4 | 1.25 | 5.82 | 4.19 |
| ENSMUSG00000006056 | Calcoco2 | 3.66 | 4.53 | 4.19 |
| ENSMUSG00000056632 | Dsg3 | 0.00 | 2.93 | 4.19 |
| ENSMUSG00000043333 | Rhbdl2 | 0.22 | 3.91 | 4.28 |
| ENSMUSG00000030838 | Ush1c | 1.15 | 5.40 | 4.38 |
| ENSMUSG00000039661 | Dusp26 | 1.15 | 4.29 | 4.38 |
| ENSMUSG00000041124 | Msantd4 | 5.83 | -1.99 | 4.38 |
| ENSMUSG00000038366 | Lasp1 | 7.93 | -2.31 | 4.55 |
| ENSMUSG00000001802 | Lrp3 | 6.00 | -4.64 | 4.77 |
| ENSMUSG00000024846 | Cst6 | 2.48 | 4.64 | 4.77 |

**Table S6:**
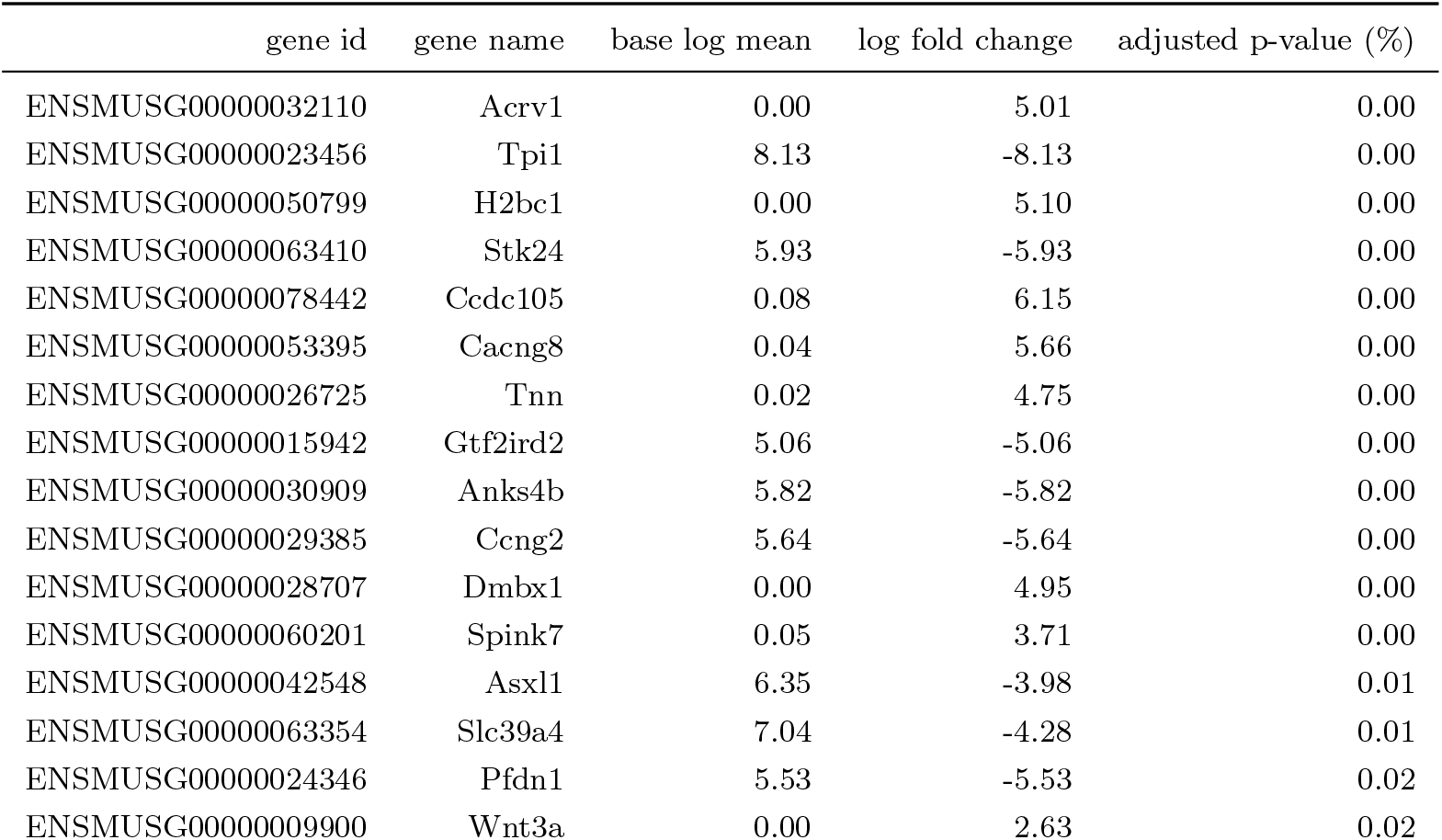
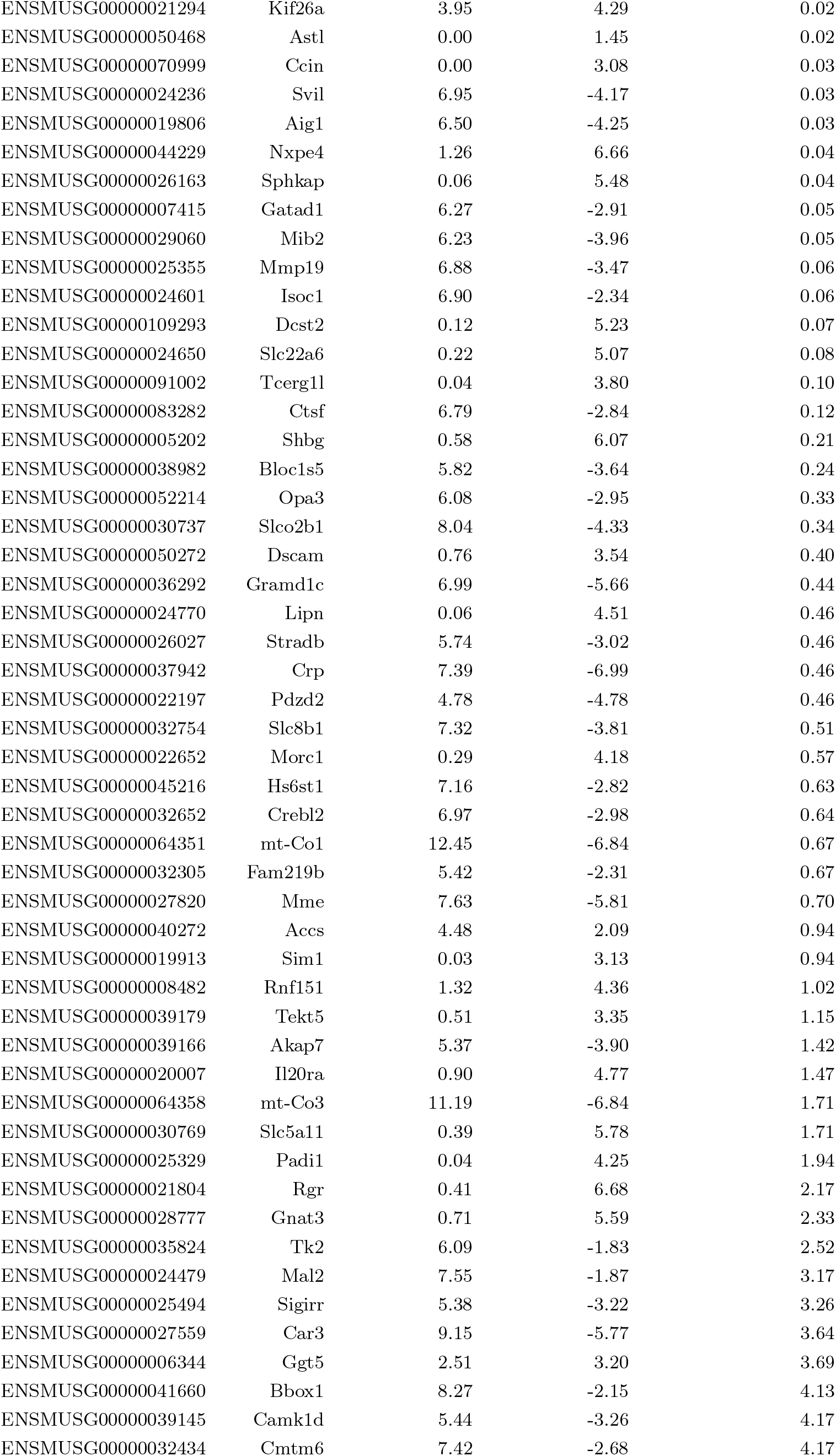
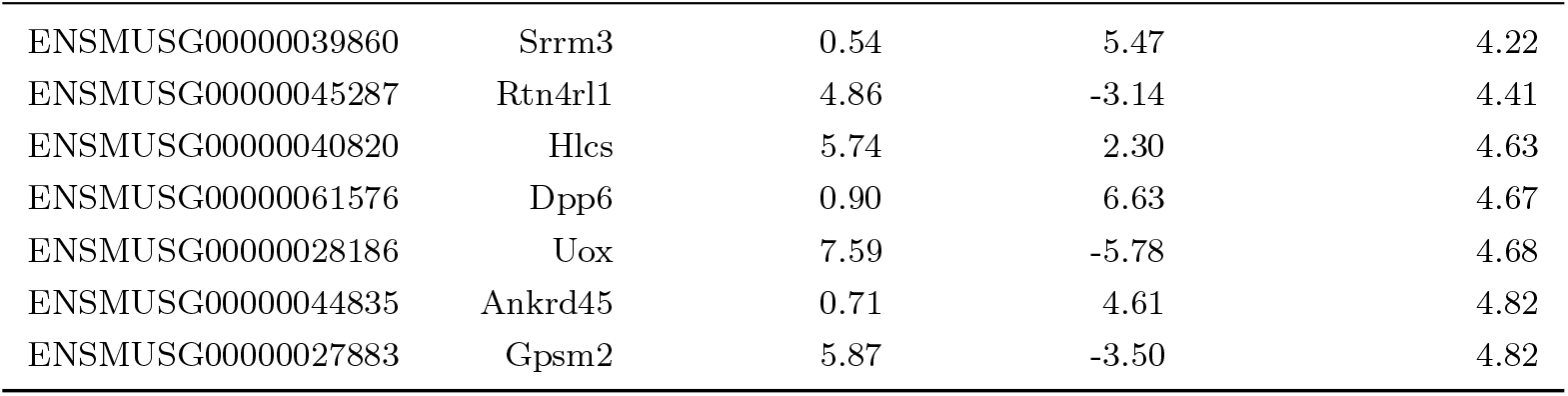
74 DE genes found by phyloDE for the Liver data with the “fdam” design.

## S6 Supplementary simulations from Stern and Crandall (2018)

### Setting

#### Crayfish Dataset

Stern and Crandall (2018) studied the mechanisms underlying vision loss in crayfishes from a dataset with 14 species. The time time-calibrated maximum likelihood phylogeny was extracted from Stern *et al*. (2017), and 3 560 orthologous gene expressions were estimated using the RNA-Seq with Expectation Maximization (RSEM, Li and Dewey, 2011). As in Bastide *et al*. (2023) and as described in the main text, we estimated the base expression level *A*_*g*,1_ and the dispersion *d*_*g*_ from the empirical data, and sampled the sequencing depth *M*_*i*_ uniformly within observed minimal and maximal values of the library size across all samples.

#### Simulation Design

We normalized the tree to unit length, and kept the original number of individuals per species (one to three, see Fig S10). We then trimmed the tree to obtain three different datasets with increasing number of samples, with a total of 4, 8 and 12 species. For each dataset, we defined two groups of equal size, according to a “block” structure, so that groups contained species with belonging to the same clade. According to Bastide *et al*. (2023), this group design is one of the most challenging for all differential expression analysis methods. The first group (gray branches, Fig S10) included species *Orconectes australis* (OA) and *Cambarus nerterius* (CN) for the smallest tree (leftmost, Fig S10), plus species *C. tenebrosus* (CT) and *C. graysoni* (CG) for the medium tree (center, Fig S10), and with additional species *O. incomptus* (OI) and *C. rusticiformis* (CR) for the larger tree (rightmost, Fig S10). The second group (black branches, Fig S10) included species *C. dubius* (CD) and *Procambarus fallax* (PF) for the smallest tree (leftmost, Fig S10), plus species *C. hamulatus* (CH) and *C. setosus* (CS) for the medium tree (center, Fig S10), and with additional species *C. cryptodytes* (CC) and *P. lucifugus* (PL) for the larger tree (rightmost, Fig S10). On the largest tree, we simulated 20 independent count RNA-Seq synthetic datasets using compcodeR, with 3 560 genes, including 250 differentially expressed genes with a minimum effect size of *e* = 3, and an underlying OU process with fixed parameters equal to *λ* = 0.7 and log(2)*/α* = 0.5 (*ρ* = 0.66). We then trimmed this dataset for the two smaller trees. The trees with transformed branch lengths according these parameters are presented Fig. S10.

#### Data Analysis

As in the main text, we applied four methods differential analysis method to each dataset, namely limma-cor, eve-ttt, phylolm (Smyth *et al*., 2005; Rohlfs and Nielsen, 2015; Ho and Ané, 2014b) and phyloDE, with the same parameters, and corrected *p*-values associated with the group effect with the same Benjamini-Hochberg procedure (Benjamini and Hochberg, 1995). We report the False Discovery Rate (FDR) and the True Discovery Rate (TPR) with a target threshold of 10%.

### Results

Results are presented Fig. S10. For all synthetic simulated dataset sizes, phyloDE had the lowest FDR, that was close to the nominal rate of 10%, and had an increasing TPR, that went up to around 60% for the largest setting (12 species). Consistent with the results of Bastide *et al*. (2023), both limma-cor and phylolm failed to control for the FDR. While the FDR of limma-cor tended to stay stable over the tree size, phylolm seemed to improve with the tree size, with a FDR going from 70% (4 species tree) to around 50% (12 species tree). This is consistent with the results of the main text, where phylolm showed a particularly poor performance on the 4 species mole rat tree. Finally, eve-ttt had a TPR that was consistently below the TPR of phyloDE, while failing to control for the FDR.

**Figure S10:**
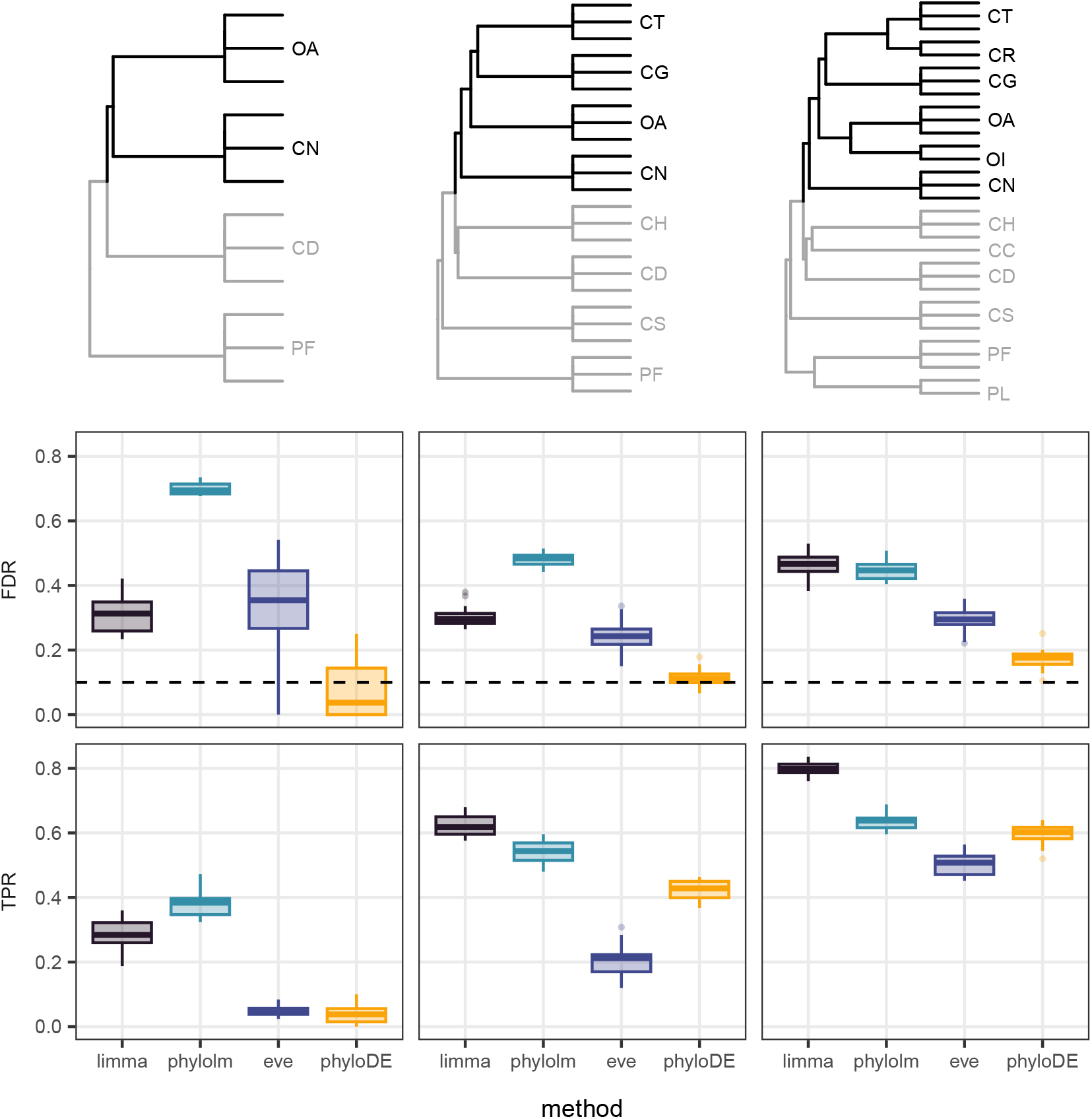
False Discovery Rate (FDR, middle row) and True Positive Rate (TPR, bottom row) at a 10% threshold (dashed horizontal line) for differential expression analysis under a “block design” scenario on subsets of the freshwater crayfish phylogenies described in Stern and Crandall (2018), with 4, 8 and 12 species, and 2 to 3 replicate per species (top row). On each tree, 20 synthetic count datasets were simulated using compcodeR following the procedure of Bastide *et al*. (2023). Trees are represented with OU-transformed trees, with parameters *λ* = 0.7 and *ρ* = 0.66. Compared methods include: limma-cor (with block correlation and empirical Bayes variance moderation with trend), eve-ttt (empirical p-values computed from 10 000 null simulations), phylolm (assuming a OU process with fixed root), and phyloDE (assuming a OU process with fixed root combined with empirical Bayes variance moderation with trend). Our method phyloDE has the lowest FDR in all settings, and shows a TPR that increases with tree size.

## S7 Supplementary simulations from Chen *et al*. (2019)

### Setting

#### Mammals Dataset

Chen *et al*. (2019) studied the gene expression evolution across 17 species and 7 tissues and detected genes with mean expression shift between one lineage and the rest of the tree. Here, we focused on the liver tissue and simulate synthetic data under two designs, isolating the rodent lineage or the primate lineage. We used a dated phylogenetic tree extracted from the larger mammals tree (Hassler *et al*., 2020). The orthologous gene expressions of 7 213 were estimated using the RNA-Seq with Expectation Maximization (RSEM, Li and Dewey, 2011). As in the previous section and as described in the main text for the molerats dataset Daunesse *et al*. (2026), we estimated the base expression level *A*_*g*,1_ and the dispersion *d*_*g*_ from the empirical data, and sampled the sequencing depth *M*_*i*_ uniformly within observed minimal and maximal values of the library size across all samples.

#### Simulation Design

We normalized the tree to unit length, and kept the original number of individuals per species (one to three, see Fig S11). We tested the lineage-specific expression shifts detection for the two lineage: primate (on the left, Fig S11) and rodent (on the right, Fig S11). On each tree, we simulated 20 independent count RNA-Seq synthetic datasets using compcodeR, with 7 213 genes, including 250 differentially expressed genes with a minimum effect size of *e* = 3, and an underlying OU process with fixed parameters equal to *λ* = 0.7 and log(2)*/α* = 0.5 (*ρ* = 0.66). The trees with transformed branch lengths according these parameters are presented Fig. S11.

#### Data Analysis

As in the main text and in the simulation study from Stern and Crandall (2018), we applied four methods differential analysis method to each dataset, namely limma-cor, eve-ttt, phylolm (Smyth *et al*., 2005; Rohlfs and Nielsen, 2015; Ho and Duchêne, 2014) and phyloDE, with the same parameters, and corrected *p−*values associated with the group effect with the same Benjamini-Hochberg procedure (Benjamini and Hochberg, 1995). We report the False Discovery Rate (FDR) and the True Discovery Rate (TPR) with a target threshold of 10%.

### Results

Results are presented Fig. S11. For all synthetic simulated design (primate or rodent), all methods had a FDR higher than the nominal rate of 10%. This is consistent with the results from the original article (Chen *et al*., 2019), where they estimated a high value of the FDR of the analysis using permutations of the original datasets. This experimental design is particularly challenging for two main reasons. First, it uses a “block design” to test for lineage-specific expression shifts, which is known to be a difficult problem (Bastide *et al*., 2023). Second, this dataset has only a few replicates per species, with only one or two replicates for 10 species out of 17 species, which can explain why the analysis is underpowered. In contrast, simulation studies from Daunesse *et al*. (2026) included 3 samples per species. For all synthetic simulated design (primate or rodent), phyloDE had the lowest FDR among the four methods (below 25%, close to the nominale rate of 10% for the rodent lineage), with a TPR around 30% higher that the TPR of eve-ttt around 25%, suggesting that this method is still competitive with eve-ttt even on larger phylogenies. Consistent with the results of the main text and on the simulation study from Stern and Crandall (2018), both limma-cor and phylolm failed to control for the FDR with values above 75%.

**Figure S11:**
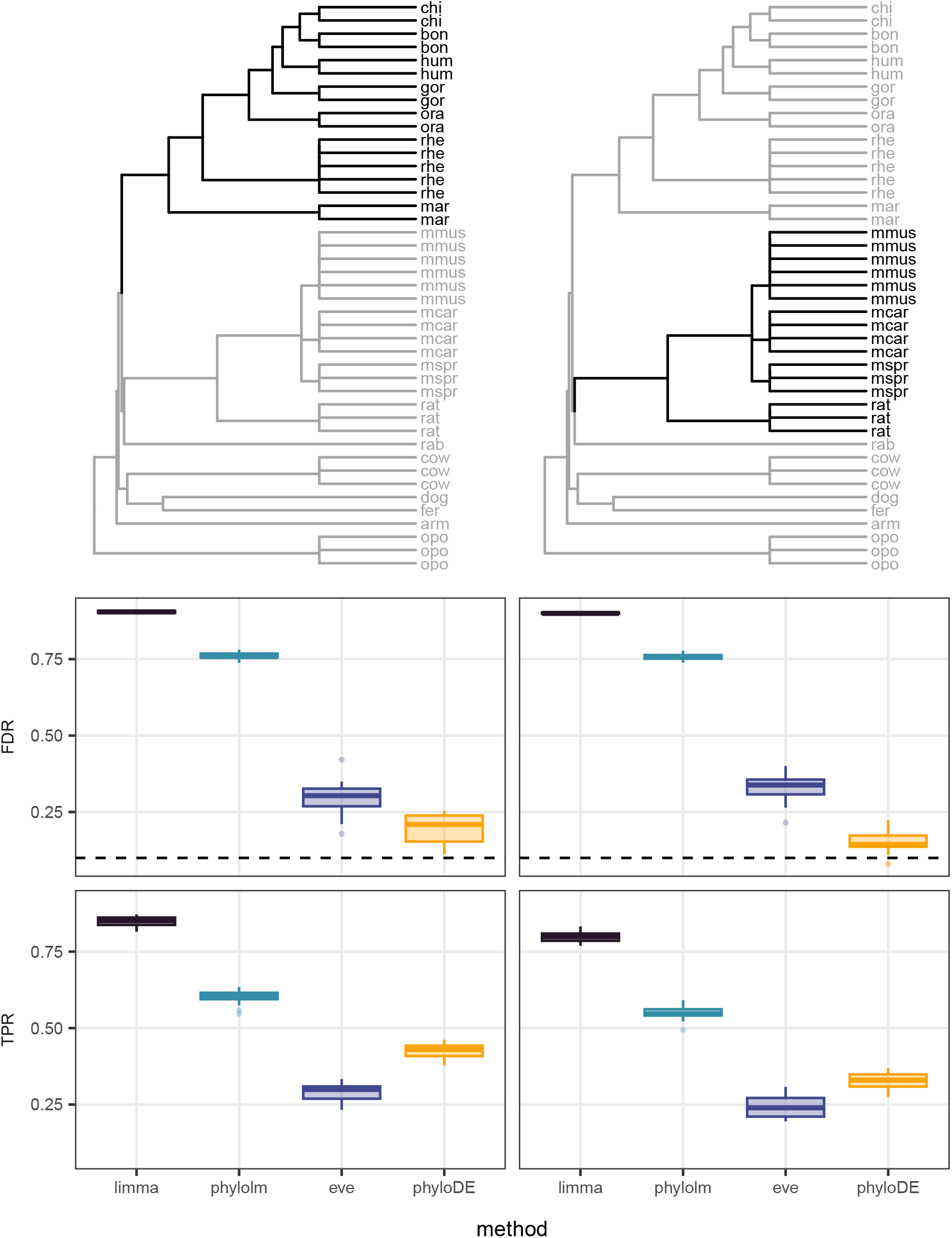
False Discovery Rate (FDR, middle row) and True Positive Rate (TPR, bottom row) at a 10% threshold (dashed horizontal line) for differential expression analysis under under two lineage-specific shift scenarios on the mammalian phylogeny from Chen *et al*. (2019) (liver tissue dataset). On each tree, 20 synthetic count datasets were simulated using compcodeR following the procedure of Bastide *et al*. (2023). Trees are represented with OU-transformed trees, with parameters *λ* = 0.7 and *ρ* = 0.66. Compared methods include: limma-cor (with block correlation and empirical Bayes variance moderation with trend), eve-ttt (empirical p-values computed from 10 000 null simulations), phylolm (assuming a OU process with fixed root), and phyloDE (assuming a OU process with fixed root combined with empirical Bayes variance moderation with trend). Our method phyloDE has the lowest FDR in all settings, and shows a TPR that increases with tree size.

